# Imaging the large-scale and cellular response to focal traumatic brain injury in mouse neocortex

**DOI:** 10.1101/2024.04.24.590835

**Authors:** Yelena Bibineyshvili, Thomas J. Vajtay, Shiva Salsabilian, Nicholas Fliss, Aastha Suvarnakar, Jennifer Fang, Shavonne Teng, Janet Alder, Laleh Najafizadeh, David J. Margolis

## Abstract

Traumatic brain injury (TBI) affects neural function at the local injury site and also at distant, connected brain areas. However, the real-time neural dynamics in response to injury and subsequent effects on sensory processing and behavior are not fully resolved, especially across a range of spatial scales. We used *in vivo* calcium imaging in awake, head-restrained male and female mice to measure large-scale and cellular resolution neuronal activation, respectively, in response to a mild/moderate TBI induced by focal controlled cortical impact (CCI) injury of the motor cortex (M1). Widefield imaging revealed an immediate CCI-induced activation at the injury site, followed by a massive slow wave of calcium signal activation that traveled across the majority of the dorsal cortex within approximately 30 s. Correspondingly, two-photon calcium imaging in primary somatosensory cortex (S1) found strong activation of neuropil and neuronal populations during the CCI-induced traveling wave. A depression of calcium signals followed the wave, during which we observed atypical activity of a sparse population of S1 neurons. Longitudinal imaging in the hours and days after CCI revealed increases in the area of whisker-evoked sensory maps at early time points, in parallel to decreases in cortical functional connectivity and behavioral measures. Neural and behavioral changes mostly recovered over hours to days in our M1-TBI model, with a more lasting decrease in the number of active S1 neurons. Our results in unanesthetized mice describe novel spatial and temporal neural adaptations that occur at cortical sites remote to a focal brain injury.

## Introduction

Traumatic brain injury (TBI) can lead to long-lasting neurological, behavioral, cognitive and emotional deficits, and represents a major public health problem with limited existing treatments. Focal TBI can cause irreversible tissue damage at the site of injury, leading to neurological dysfunction. Secondary processes often distant from the injury site are also thought to be a major contributor to TBI-related deficits (Carron et al., 2016; Johnstone et al., 2014; Meaney et al., 2014). How focal injury triggers long-lasting and widespread secondary processes is an important area of TBI research.

Focal TBI in cortical impact and contusion models involves disruption of neural cell membranes and rapid imbalance of ionic concentrations (e.g., Na+, K+, Ca2+, Mg2+) important for cellular excitability (Pevzner et al., 2017). The rise in extracellular K+ strongly depolarizes neurons, leading to acute release of neurotransmitters (e.g., glutamate, acetylcholine, GABA) (Pevzner et al., 2017), activation of postsynaptic receptors, and pathophysiological spreading cellular activation, called cortical spreading depolarization (CSD). CSD is a slow wave of massive neuronal depolarization, initiated at the injury site and propagated by K+ diffusion, followed by temporary suppression of neural activity (Enger et al., 2015) and is considered a key event following TBI (Bogdanov et al., 2016; Lauritzen et al., 2011; Charles and Brennan, 2009; Dreier et al., 2011; Hartings et al., 2009; 2011; 2017; Fabricius et al., 2005). Because of the suppression, CSD has also been termed cortical spreading depression (Dreier, 2011; Lindquist and Shuttleworth, 2012; Chang et al., 2010; Sawant-Pokam et al., 2017).

TBI-induced changes in brain function occur over extended spatial and temporal scales, with effects spanning the molecular, synaptic, cellular, and network levels (Alwis et al., 2013; Gaetz, 2004; Greve and Zink, 2009; McIntosh et al., 1996; Cantu et al., 2014; Hunt et al., 2010). Functional changes are complex and can vary across cell types and brain regions, but commonly include decreased neuronal excitability and depressed excitatory synaptic function (Goforth et al., 2011; Li et al., 2014; Sawant-Pokam et al., 2020) starting within minutes of injury and lasting days. Altered cortical synaptic plasticity, including the capacity for long-term potentiation, can persist for weeks (Sanders et al., 2000), depending on the severity of injury, and can be similar to the time scale of behavioral and memory deficits (Tucker et al., 2016; Washington et al., 2012).

For example, Ding et al. (2011) found that motor cortex compression injury induced an initial reduction in sensory-evoked spiking responses in the somatosensory cortex that recovered gradually over 1 h, then became hyperexcitable. Similarly, neurons near (1-2 mm) a motor cortex controlled cortical impact (CCI) injury site showed initial depression followed by hyperexcitability that lasted up to 14 d (Ping and Jin, 2016). TBI-related changes in brain functional connectivity, measured with quantitative network analysis, have also been reported (Aerts et al., 2016). Using measures such as nodal degree, connectivity strength, global efficiency, clustering coefficient, modularity and centrality (Alhourani et al., 2016; Nakamura et al., 2009), network analysis has found TBI-induced reductions in functional connectivity between brain regions across mild to severe TBI (Pandit et al., 2013; Han et al., 2016). Functional connectivity is disrupted by damage to white matter tracts or diffuse axonal injury (Kinnunen et al., 2011), and in brain areas remote from the site of focal TBI (Palacios et al., 2017; Stephens, et al., 2018; Sharp et al., 2014).

The neural response to TBI both at focal and distal sites, including spatial and temporal aspects of the spread of neural activity from the injury site to distal brain areas and the capacity of specific circuits for recovery, require further investigation (Hosseini-Zare et al., 2017; Alwis et al., 2013). Because of the difficulty in monitoring brain activity during TBI induction (compared to select post-TBI time points), there is a gap in experimental data on the brain’s response to TBI at the moment of injury. Our goal was to capture the immediate neural response to TBI across broad cortical areas and local cellular populations, and to relate the injury response to subsequent recovery of function.

## Methods

### Ethical approval

All experiments were approved by Rutgers University Institutional Animal Care and Use Committee (IACUC; protocol 13–033) and in accordance with the United States federal guidelines of the Animal Welfare Act. All personnel were trained and approved to work with laboratory animals in order to minimize animals’ pain and suffering. The authors understand the ethical principles related to scientific publishing and affirm that the research reported here complies with the journal’s animal ethics checklist.

### Experimental overview

This study aimed to investigate the effects of a focal controlled cortical impact (CCI) injury of primary motor cortex (M1) on mesoscopic and cellular cortical activity and behavior. Mice were implanted with a cranial window and head post (see Surgical procedure, below) 10 to 14 days before the day of M1-CC1 injury. In the 7 days before injury, mice were habituated to head-fixation on a crossbar and holder for positioning under the microscope, as in previous studies (Park et al., 2016; Lee and Margolis, 2016). Experimental sessions for widefield calcium imaging consisted of measuring spontaneous activity (8 trials, 20 s each) followed by measuring sensory-evoked activity (30 trials, 5 s each). Widefield imaging sessions were followed by 10 min behavior observations in the home cage, including the number of grooming and rearing events. In two-photon imaging sessions, spontaneous activity was measured in a single trial of at least 200 s; sensory-evoked activity measured in at least 30 trials of 5 s duration. No sensory-evoked activity was recorded on the day of injury (session 5) in two-photon experiments. As summarized in the timeline of **figure 1E**, three baseline sessions were performed before the day of injury. On the day of injury, one session was performed before injury followed by measures during CCI application. Additional sessions were performed at 20 and 60 min post-CCI, followed by sessions on days 1, 3, 7 and 14 post-CCI. For widefield imaging there was an additional session at 8 weeks post-CCI. On the day of injury mice were randomly divided into Injury or Sham groups. A small craniotomy (approximately 0.6 - 1.0 mm) was made over the left hemisphere of M1 under isoflurane anesthesia in both Injury and Sham groups. Imaging sessions on the day of injury were started 1 h after the cessation of anesthesia. Mice were awake and head-restrained under the microscope for all recordings.

**Figure 1.**
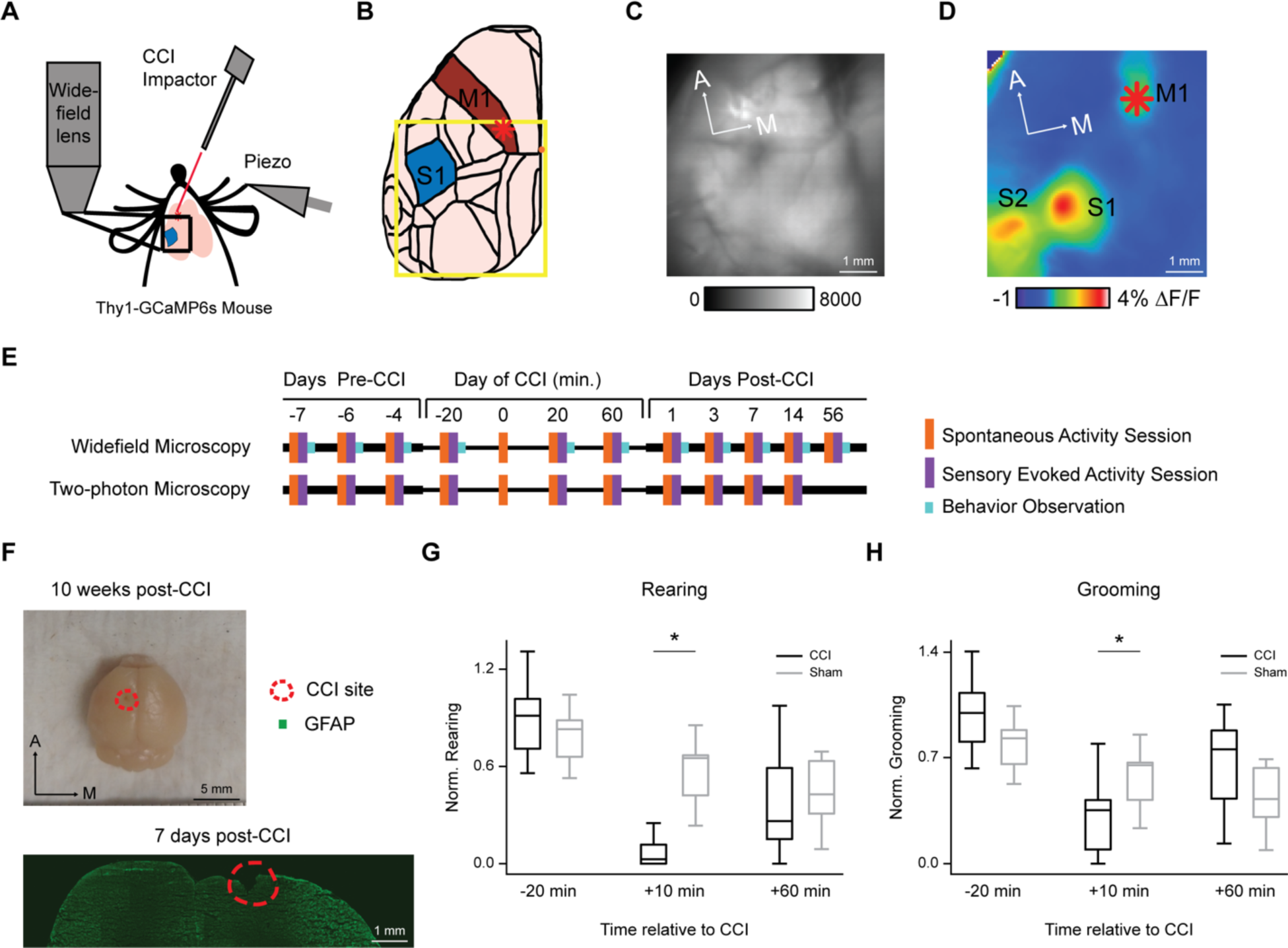
Experimental overview. **A,** Schematic of experimental setup for imaging cortical calcium signals in transgenic reporter mice during focal controlled cortical impact (CCI). CCI probe velocity was 4 m/s and 0.5 mm diameter. Piezoelectric actuator (Piezo) provided 0.5 m/s velocity, 0.1 Hz frequency whisker deflection. Blue square indicates field of view for widefield imaging. **B,** Schematic of cortical areas within the field of view of the left hemisphere of mouse brain (based on Allen Mouse Brain Common Coordinate Framework; https://scalablebrainatlas.incf.org/mouse/ABA_v3). M1, primary motor cortex (red); S1, primary somatosensory cortex (blue). Red asterisk indicates M1 injury site, where CCI was applied. Orange dot indicates Bregma. **C,** Example widefield fluorescence image of cortical surface through the transparent skull preparation. **D,** Example sensory-evoked cortical activation map, as used to localize CCI to M1. Map shows pseudocolored change in calcium signal fluorescence (% ΔF/F) following D2 whisker stimulation averaged for multiple trials. Field of view and subject are the same in C and D. S2, secondary somatosensory cortex. Red asterisk shows CCI injury site. **E,** Timeline of imaging experiments. Colors indicate sessions of recording spontaneous activity (orange), sensory-evoked activity (purple), and behavioral observations (cyan). Separate groups of mice were used in widefield and two-photon experiments. “0 m” indicates the time of CCI on the day of injury. **F,** Top, Dorsal view brain extracted 10 weeks post-CCI. Bottom, Coronal section stained for GFAP (astrocytes) and DAPI (nuclei). CCI-induced tissue damage outlined by red dashed line. **G,H,** Behavioral effects of M1 CCI. Left, mean normalized number of rearing events (left) and grooming events (right) for Injury (n=12) and Sham groups (n=11), measured 20 min before, 0 min, and 60 min after the M1-CCI imaging session.

### Animals

Male and female Thy1-GCaMP6s calcium reporter mice (GP4.3; heterozygous; purchased from Jackson Labs, Jax stock number 024275) (Dana et al., 2014) were bred and raised in sterile conditions with 12-hours dark/light cycle and free access to food and water (temperature 23– 25°C, humidity <10%). Mice were 9-14 weeks old on the day of injury, or up to 36 weeks old in a few cases. Three weeks before injury, after surgical implant of the imaging window and head post, mice were transferred to a reversed light cycle room. Subsequent handling and experiments were performed in the dark phase of the diurnal cycle. After recovery from surgery, mice were gradually accustomed to head fixation, first by sitting on the hands of the experimenter and then by undergoing gentle manually applied head restraint, before the head post was affixed to the holder used for imaging experiments. Mice sat inside a plexiglass tube attached to the holder.

The above behavioral handling was performed 3-4 times per day over approximately 10 days before imaging experiments began. After imaging sessions started, the health of subjects was monitored before every experimental session. Mice were removed from the study if any signs of adverse health were present (2/27 mice in widefield group). In the two-photon group, 7/22 mice were removed from the study because of insufficient optical quality of the cranial window. An additional 7/22 mice were removed from the study following application of the CCI because of cortical bleeding or bruise formation, affecting optical quality of the cranial window. Mice removed from the study for these reasons were killed by carbon dioxide euthanasia and cervical dislocation as a secondary method. If brains were to be harvested for histology, mice were killed by ketamine/xylazine anesthesia and transcardial perfusion, as below in the section “Histological measurement of CCI severity”.

### Surgical procedure

Mice were anesthetized with isoflurane (4% induction, 0.8–1.5% maintenance) and placed on a stereotaxic frame, where an animal’s body temperature was maintained 36°C with a feedback-controlled heating blanket. After subcutaneous lidocaine injection of the scalp, the skull was exposed and cleaned for application of bonding agent (iBond, Heraeus Kulzer, Hanau, Germany). For transparent transcranial windows, one layer of light-curable dental cement (Tetric Evoflow, Ivoclar Vivadent, Schaan, Lichtenstein) was applied. A second type of light-curable dental cement (Charisma, Heraeus Kulzer, Hanau, Germany) was applied at the border of the first layer of cement and the skin, to form a chamber for immersion fluid during imaging experiments. For two-photon imaging, after applying bonding agent and dental cement, a 4 mm diameter craniotomy was performed above the somatosensory cortex by careful use of a dental drill (Osada EXL-M40, Los Angeles, CA). A 3 or 4 mm round glass coverslip was placed on the dura mater and fixed in place with Tetric Evoflow dental cement surrounding the edge. An aluminum head post was glued and cemented to the right side of a skull. Rimadyl was administered subcutaneously. Mice recovered from surgery with free access to food and water in a reversed light cycle room (12-hours dark/light) and were monitored daily.

### Controlled cortical impact

On the day of injury, mice were anesthetized with isoflurane as in the original surgery (4% induction, 0.8–1.5% maintenance), and an approximately 1 mm craniotomy made over the left motor cortex (M1), leaving the dura intact. M1 location was targeted from cortical mapping during the baseline imaging sessions; approximate coordinates were 1 mm anterior, 0.6 mm lateral from Bregma. A small amount of Charisma dental cement was placed posterior to the craniotomy as a barrier between the injury site and the imaged regions of cortex. The craniotomy was temporarily closed with parafilm and the mouse returned to the home cage for 1 h. After 1 h recovery, the mouse was placed under the microscope for a standard imaging session. The mouse was returned to the home cage again for 10 min for pre-CCI behavior observation and then back to the microscope for recording of spontaneous activity during M1-CCI. Mice were randomly assigned to Injury or Sham groups. Parafilm was removed from above the craniotomy and the CCI probe positioned over M1 in Injury mice or approximately 2 cm to the side of the head for Sham mice. During a spontaneous activity movie, the CCI device was activated, initiating M1 injury (in Injury mice).

To perform M1-CCI during imaging, the probe of the device (Hatteras instruments, PinPoint, Precision cortical impactor, Model PCI3000, Cary, NC) was positioned at a 42 ± 5° angle. This allowed the probe to fit under the microscope allowing for simultaneous imaging. CCI was delivered with the parameters: velocity, 4 m/s; duration, 85 ms, penetration depth 1.8 ± 0.2 mm. The probe diameter was 0.5 mm to fit within the 1 mm craniotomy. The relatively small probe size and parameters used for CCI produced a mild or moderate, rather than severe, TBI. After the imaging session, bleeding (if present) was absorbed and the craniotomy closed with Evoflow dental cement. Post-CCI (Post-Sham) behavior was observed for 10 min in the home cage, and then imaging was repeated at 20 min and 60 min.

### Histological measurement of CCI severity

To characterize the injury severity histologically, brain tissue loss at the injury site was measured at 7 d post-CCI in a subset of wildtype mice (n = 4). Mice were surgically prepared, recovered, and CCI delivered in the same way as described above for mice in imaging groups. After the M1-CCI, the craniotomy was sealed with dental cement and mice returned to the home cage for 7 d. After 7 d, mice were deeply anesthetized (120 mg/kg ketamine plus 24 mg/kg xylazine, intraperitoneally) and transcardially perfused with 0.1 M phosphate buffer followed by 4% paraformaldehyde (PFA) in 0.1 M phosphate buffer. Brains were postfixed for 24 h in 4% PFA, followed by 24 h cryoprotection in 30% sucrose solution and 0.1 M of phosphate buffer. After cryoprotection, each brain was placed in PBS solution with sodium azide (0.02 %). Frozen sections (20 µm;) were prepared throughout the site of injury on the cortex in a 1:20 series. DAPI (1:1000, Sigma) and GFAP staining (1:500 GFAP antibody MAB3402, Millipore with following Alexa Fluor goat anti-mouse 488 Invitrogen 1:250) was used to label nuclei and staining astrocytes. Injury site was visualized with a Zeiss Axio Observer Z1 inverted fluorescence microscope, imaged at 10x or 20x and exported to tiff using Zeiss LSM 800 Confocal software, using the mosaic tile function. The area of the injury site was visually determined in ImageJ (2-3 images per mouse). The volume of tissue loss was calculated under the assumption of a circular perimeter of the injury site.

### Widefield calcium imaging

Widefield (mesoscopic) calcium imaging was performed in awake, head-fixed Thy1-GCaMP6s mice using a custom macroscope, as previously described (Minderer et al., 2012; Steinmetz et al., 2017; Zhu et al., 2018). Fluorescence excitation was provided by a high-power 60-chip 435 nm LED (±15 nm half-width; Roithner Lasertechnik), with intensity controlled by a stabilized current driver (Prizmatix) and filtered through a 479 nm/40 nm excitation filter (Chroma). Excitation light was reflected by a 50 mm dichroic mirror (Q470lp, Chroma) and focused through a 25 mm f0.95 lens (Navitar D-1795 or D-2595; size of each pixel was 33 or 49 μm width depending on magnification). A MiCam Ultima-L (Brain Vision, Inc.) camera system with 100 x 100 pixel CMOS sensor was used for imaging. Fluorescence emission was collected by a 50 mm f0.95 lens (D-5095; Navitar), filtered (535 nm/40 nm; Chroma) and reflected to the camera. Acquisition and length of movies were controlled by the MiCam Ultima imaging system software (Brain Vision, Inc.). Frame rate of acquisition was 100 Hz. The maximal movie duration at this frame rate was 20 s. For longer events, multiple movies were acquired with a 13 s inter-movie interval, and the gap between consecutive movies was interpolated offline. Importantly, data analyses were performed on the non-interpolated data, so interpolation did not affect the interpretation of the data.

### Two-photon laser-scanning microscopy

Chronic two-photon calcium imaging was performed in awake, head-fixed Thy1-GCaMP6s mice, similar to previously described (Margolis et al., 2012), using a Neurolabware microscope (Los Angeles, CA). Fluorescence excitation was provided by a Ti:sapphire laser system (Mai Tai HP, Newport Spectra Physics, Santa Clara, CA; 100 fs laser pulses; 920 nm wavelength) and scanned across the sample with resonant-galvanometer scanning mirrors (models CRS 8K and 6210, Cambridge Technology, Bedford, MA). A Pockels cell (Conoptics, Danbury, CT) was used for laser intensity modulation, and a triggerable shutter (Uniblitz VCM-D1, Vincent Associates, Rochester, NY) blocked the beam when not acquiring. Fluorescence excitation was collected through a Nikon (Tokyo, Japan) 16x/0.8 NA water immersion objective. Photomultiplier tubes (PMTs; Hamamatsu, Japan) were used for fluorescence detection. Note that PMT gain was optimized for physiological cellular signals and was not adjusted during the large-amplitude, CCI-induced Wave. This resulted in PMT saturation during the CCI-induced Wave. The motivation for this was to collect cellular signals before, during the onset, and after the Wave; however, PMT saturation precluded analysis of detailed Wave properties in two-photon experiments. Effective frame rate was 15.46 images/s with 540×720 pixel resolution. Acquisition was controlled by Neurolabware Scanbox software.

### Spontaneous activity and sensory stimulation

During widefield imaging sessions, spontaneous and sensory-evoked cortical activity was measured. Spontaneous activity was recorded in 8 movies 20 s duration with 13 s inter-movie interval. After that, the D2 whisker was connected to the piezo bending element (approximately 5 mm from the whisker base) using a drop of cyanoacrylate glue. Stimulation trials activated the piezo bending element, causing mechanical deflection of the whisker by 0.3-0.5 mm (approximately 14 degrees) in 30 ms. One whisker deflection was delivered per trial, 1 s after trial onset, with 8 s inter-trial interval. Imaging frame rate was 100 Hz. 30 trials of sensory-evoked responses were acquired per session. For two-photon imaging sessions, at least 30 trials of D2 whisker stimulation were delivered in the same way as for widefield imaging. The stimulation timing was acquired during the 200 s duration two-photon movie for offline synchronization.

### Data processing and analysis

Widefield fluorescence data were imported into ImageJ (http://rsbweb.nih.gov/ij/) as 16-bit images, converted to tiff format, and calcium signal fluorescence changes (% ΔF/F) determined using custom routines. ΔF/F was calculated pixel-wise by subtracting the baseline (F; the average of the first 30-200 frames) from each frame in the video and dividing each frame by the same baseline.

For spontaneous activity measures, including movies acquired during the CCI, all 8 (20 s duration) movies were concatenated before ΔF/F calculation. Timeseries data were extracted from ROIs defined in ImageJ and imported into Matlab (Mathworks). A grid of ROIs covering the field of view (each 10 by 10 pixels, 0.51 mm^2^), or a single ROI in the D2 barrel column of S1, were used to analyze timeseries data for features of the CCI-induced calcium signals that we termed “initial response” (IR), “Wave”, “Depression” and “Recovery”. ROIs were excluded from further analysis if they covered dental cement or areas of cortex obscured by bruise or bleeding. The 13 s inter-movie interval created a blank period between the successive 20 s movies when no data were recorded. The blank period was interpolated in the figures for display. For signal amplitude analysis, ROIs were included that showed maxima or minima during recorded periods, and excluded if they had a presumed peak within a blank period.

Peak IR and Wave signal amplitudes were measured from ΔF/F timeseries data as the mean of 7 or 51 frames around the maximum, respectively, to account for any fast signal fluctuations. The Depression, or signal decrease below the baseline level, was measured as the mean of 51 frames around the signal minimum within 66 s (within two movies) after the peak Wave signal. The Recovery, or return of calcium signal fluorescence to baseline following the Wave, was measured as the mean of 51 frames around the signal maximum within 54 s after the Depression. Recovery was variable in different cortical regions; only ROIs that showed Recovery within 160 s (duration of 8 movies) were included in Recovery analysis. Group means were calculated for each ROI or averaged across ROIs.

For analysis of spatial distributions, circular ROIs (7 pixel diameter; 0.5 mm; 38.5 mm^2^) were placed approximately 2.2 mm from the M1 injury site, spaced 0.7 mm apart, along directions 22.5^∘^, 45^∘^ and 67.5^∘^ relative to the midline in the posterior-lateral direction. For analysis of calcium signal area, masked movies were created by binarizing pixels that exceeded an amplitude threshold (> 20% maximum amplitude for IR and Wave; < 20% minimum for Depression; > 10% maximum amplitude for Recovery, relative to Depression). Pixels within 10 pixels of the edge of the field of view were excluded to avoid edge effects. The sum masked pixels in each frame represented the signal area dynamics. Area values were reported as the average value of peak ±3 frames for IR, and ±25 frames for Wave, Depression or Recovery in units of mm^2^. Trajectories of CCI-evoked signals were analyzed in Matlab. For the Wave, the position of the maximum signal was measured every 1s (100 frames) within the two movies following the IR. Pixels within 10 pixels of the edge of the frame were excluded to avoid edge effects. Trajectories of Depression were analyzed in the same way but for the signal minimum. Trajectories and M1 injury sites were plotted in Adobe Illustrator on images registered to the midline for comparison across subjects.

For sensory-evoked calcium signal analysis, ΔF/F was calculated separately for each of 30, 5 s movies. To estimate signal onset location, all movies were spatially averaged and the location of maximal intensity located in the 10 frames (100 ms) following whisker deflection. To measure peak amplitude, timeseries data was extracted from a 10×10 pixel rectangular ROI placed over the signal onset location. Peak amplitude was defined as the difference between the mean of 30-100 pre-stimulus baseline frames and the maximal signal up to 150 ms after stimulus. To calculate the spatial area of sensory-evoked signals, 5 frames around the time of the peak signal were averaged. Area was measured as the summation of pixels exceeding 50% peak amplitude and converted to mm^2^.

Two-photon microscopy data were analyzed using Scanbox Yeti routines in Matlab and CaImAn (Giovannucci et al., 2019) and Suite2P routines in Python. CaImAn was used for alignment, ROI definition and signal extraction from individual ROIs. Both Yeti and CaImAn were used to generate maps of active neurons (Yeti used coactive pixels within the first 300 frames of each trial; CaImAn used full trial data). Final determination of ROI placement and size was verified manually. The total number of neurons per field of view was estimated by manual inspection. The Wave amplitude was truncated in two-photon recordings because of PMT detector saturation. The number of neurons active during the Wave is a lower estimate due to the brightness of cellular and neuropil fluorescence that overwhelmed individual neurons. Depression amplitude was calculated as described for widefield imaging. For analysis of spontaneous activity, events were defined as calcium transients with amplitude greater than 4x the baseline SD. For analysis of sensory-evoked activity, active/responsive cells were defined as detected by Suite2P. The amplitude and duration (full width at half maximum; FWHM) of trial-averaged calcium signals following whisker deflection were calculated for each responsive cell.

### Functional connectivity analysis

Cortical networks are often defined in neuroimaging studies as regions (or nodes) with covarying activity, or functional connectivity. To analyze functional connectivity of cortical networks, timeseries data from movies of spontaneous activity were extracted from 5×5 pixel ROIs (as in Fig.7A) that were considered as the nodes for construction of functional networks. Individual trials were considered outliers and excluded from further analysis if the S.D. of the timeseries was 2x the average S.D. of other trials in that session. Considering the fluorescence signal of each ROI for the duration of *T* timepoints during the recording session, we obtain timeseries *s_i_*(*t*) for *i* = 1, …, *N* and *t* = 1, …, *T*.

The undirected weighted functional connectivity networks were then constructed for each trial of the recording by computing their adjacency matrices. The weighted network analysis, in contrast to the binary one, makes it possible to track changes in the strength of the connections following injury. To construct the network in each trial, wavelet transform coherence (WTC) (Addison et al., 2002) for all 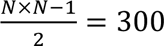 possible ROI pairs were computed. WTC enables determination of the coherence between two timeseries as a function of time and frequency. WTC decomposes a single timeseries into time and frequency space by consecutively convolving the time series with scaled and translated versions of a wavelet function *Ψ*_0_(*t*) (here we used Morlet). For a timeseries *s_i_*(*t*) of length *T*, its continuous wavelet transform, *W_si_*(*t*, *n*), sampled at equal time steps of size *Δt* is obtained as

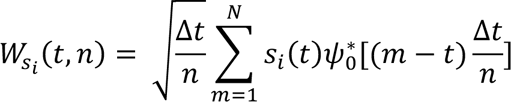

where the parameter *n* represents the wavelet scale. The WTC 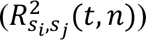 of two time series *s_i_*(*t*) and *s*_-_(*t*) is defined as

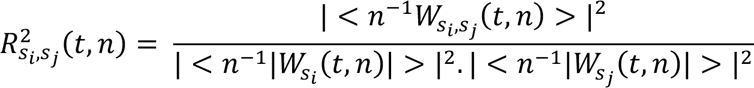

where <. > represents smoothing in both time and scale. Computing the WTC between all ROI pairs, connectivity strength between each ROI pairs was defined by averaging the WTC values across the frequency range of 1 − 5 Hz and the time duration *T* of each trial. The frequency range of 1 − 5 Hz was selected based on the results of previous GCaMP mice studies, showing that this frequency range is the dominant frequency band (as distinguished from slower 0.1 to 1 Hz signals) associated with the cortical activity in GCaMP mice (Vanni et al., 2017; Wright et al., 2017). Computing the connectivity for all 300 ROI pairs resulted in a series of brain network adjacency matrices of *A*_25×25_.

Data were included from mice in Injury (n=13; 7 male, 6 female) and Sham (n=12; 7 male, 5 female) groups from 10 sessions over 7 d pre-CCI to 14 d post-CCI.

### Statistical analysis

Group data are presented as mean ± SD. Statistics were calculated using Matlab or the lme4 R package. Data were analyzed and compared using paired or unpaired t-tests, one-way analysis of variance (ANOVA), two-way ANOVA, or linear mixed effects models (LME). ANOVA tests were followed by post hoc multiple comparison tests with Holm correction for multiplicity. In all cases, differences were considered significant at p < 0.05.

## Results

CSD is considered an important component of the brain’s response to injury, but there is limited data relating its spatiotemporal features to other injury-induced neural and behavioral changes. We sought to measure the widespread effects of focal TBI, and to follow the consequences of these effects over time in a transgenic mouse that expresses the genetically encoded calcium indicator (GECI) GCaMP6s throughout excitatory neurons of the cerebral cortex (Dana et al., 2014). CCI is considered a clinically relevant TBI model as it produces histopathologic and vascular changes similar to TBIs in humans (Lighthall, 1988). We chose a model of motor cortex (M1)-CCI because M1 is a clinically relevant brain area affected in TBI, and because M1 is spatially separated enough from sensory cortex (S1) to allow for investigation of S1 as a remote, connected brain area amenable to imaging. S1 is relevant because S1 in rodents and humans appears to be particularly vulnerable for induction of CSD (Lauritzen et al., 2011; Bogdanov et al., 2016) and sensory processing is commonly affected in TBI (Thielen et al., 2023). Performing in vivo calcium imaging experiments in awake mice avoids effects of anesthesia on neural activity and TBI outcome (Statler et al., 2006). The experimental preparation allowed us to resolve the immediate effects of M1-CCI and relate the effects to the recovery of spontaneous and sensory-evoked cortical activity, on the levels of cortical areas and neuronal populations, over the following weeks in the same subjects.

### Widefield calcium imaging of cortical activity during focal injury of M1

To determine the large-scale (or mesoscopic) cortical response to focal TBI, we performed *in vivo* widefield calcium imaging in awake, head-restrained Thy1-GCaMP6s transgenic reporter mice during application of a controlled cortical impact (CCI) to the whisker-related motor cortex (M1) (**Fig. 1A**). The field of view contained the M1 injury site at the anterior edge, somatosensory regions, including primary somatosensory (S1) barrel cortex, and portions of visual cortex at the posterior edge (**Fig. 1B**). Functional maps of S1 barrel cortex and M1 were obtained in order to localize the site of CCI to the M1 region on the day of injury (**Fig. 1C, 1D**). The whisker-evoked activation of S1 and downstream cortical areas such as secondary somatosensory cortex (S2) and M1 is consistent with previous widefield imaging studies in awake, head-restrained mice (Ferezou et al., 2007). We performed repeated widefield or (in separate mice) two-photon calcium imaging across multiple days before and after injury, allowing for longitudinal tracking of injury effects within each subject. The overall experimental timeline is shown in **Fig. 1E**. Imaging of both spontaneous activity and sensory-evoked activity was performed in each session, as well as post-session home cage behavioral observations in the widefield group.

To characterize the severity of TBI induced by our CCI paradigm (see Methods for device parameters), we examined the extent of tissue damage in post-mortem brain tissue. Histological analysis of the injury site indicated an injury size of 0.45 ± 0.22 mm^3^ (mean ± SD, n = 4 mice), which is relatively small compared to previous studies (Radomski et al., 2013; Hemerka et al., 2012; Tucker et al., 2016) (**Fig. 1F**). To examine the basic behavioral effects of TBI, we scored post-injury rearing and grooming when mice were returned to the home cage. Mice in the injury group showed stronger post-injury reductions in rearing (92%) and grooming (64%) compared to mice in the Sham group (rearing 44%, grooming 5%) that lasted for up to one hour (**Fig. 1G**; n = 12 mice, injury; n = 11 mice, Sham; see also Fig. 7 for the full time course of behavioral effects) [ANOVA of Linear mixed effects (LME) model (∼ Condition * Time + (1|Mouse)) showed significant interaction with condition and time for rearing (F=3.51, p=0.00015) and grooming (F=2.28, p=0.012); post hoc analysis with Holm corrected multiple pairwise comparisons showed significant differences between injured and sham, * p < 0.05. (t = −4.10, −3.95; p = 0.012, 0.028) for normalized rearing and grooming respectively at T = +10 min]. The restricted histological extent, parameters of the CCI including a relatively small 0.5 mm probe size, and short-lasting behavioral effects indicate that our CCI paradigm induced mild or moderate, not severe, TBI (Siebold et al., 2018; Chen et al., 2014; Washington et al., 2012).

To image cortical activity dynamics during TBI induction, we angled the piston of the CCI device at approximately 40 degrees for positioning under the widefield or two-photon microscope. This allowed us to image the immediate effects of TBI on cortical activity dynamics, which has been difficult to achieve experimentally. Widefield calcium imaging during CCI induction revealed a striking large amplitude, widespread cortical activation (**Fig. 2**). Upon impact, the CCI probe elicited an initial response (IR) to mechanical contact (**Fig. 2B, left**), followed by a massive traveling “Wave” within a few seconds delay after the CCI probe had left the tissue (**Fig. 2B**; note longer time scale). The Wave of increased calcium signal fluorescence progressed slowly over tens of seconds and over several millimeters of cortex in the posterior and mediolateral directions from the injury site (**Fig. 2C**) (anterior spread was not resolved because this was outside of the field of view). After the brief IR and prolonged Wave, there was a reduction of calcium signal fluorescence below the resting level that we termed a “Depression” (**Fig. 2B, 2C insets**). The Depression showed a faster phase when fluorescence decreased below the pre-CCI baseline level, followed by a slower phase when fluorescence either stayed at the same level, continued to decrease, or in some cases showed recovery within the subsequent minutes (**Fig. 2B**). Note that the Depression and Recovery responses are shown on a compressed scale in the main panels and insets because their amplitude was much smaller than the Wave.

**Figure 2.**
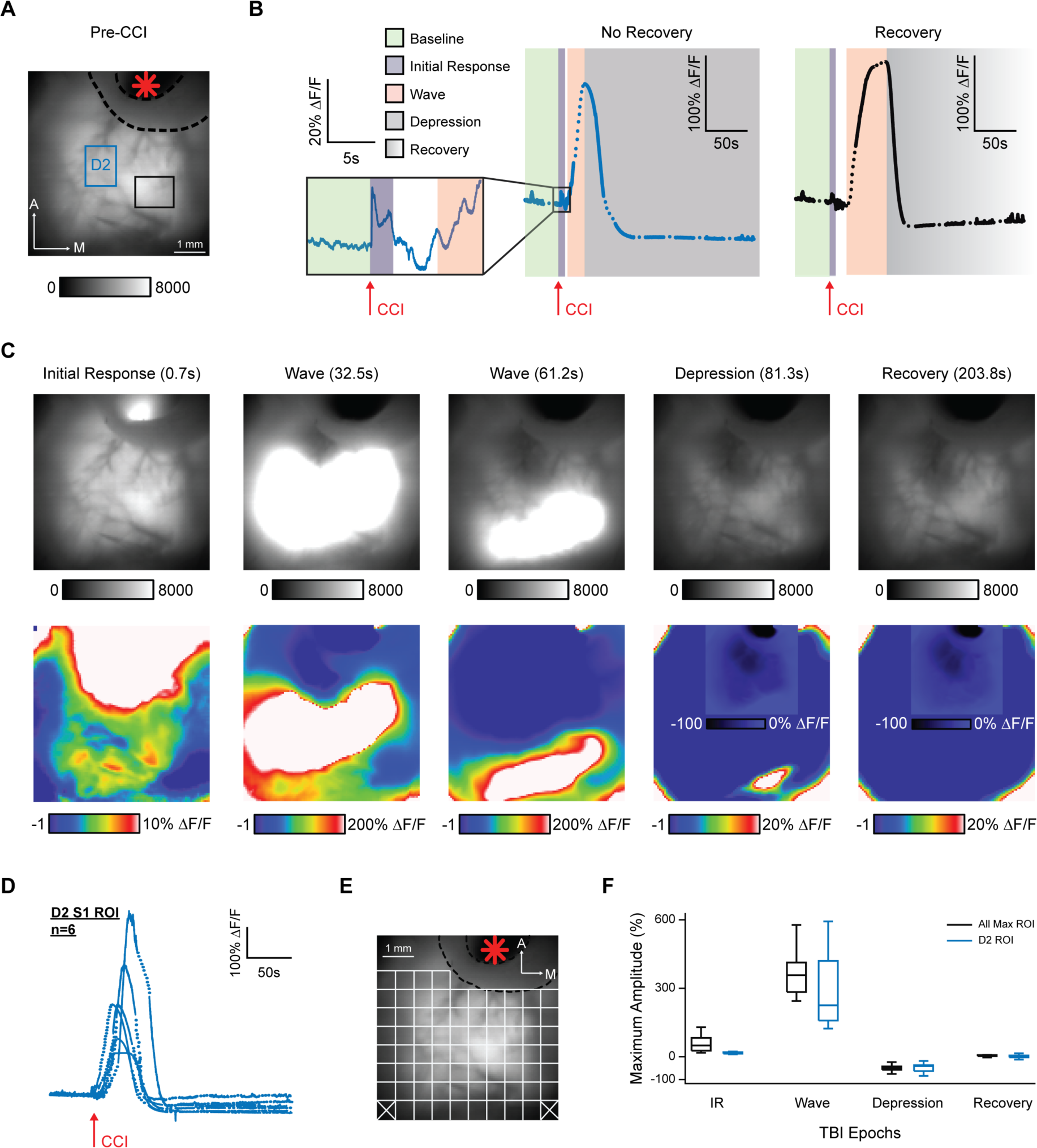
Widefield calcium imaging of cortical activation during M1-CCI focal injury. **A,** Widefield fluorescence image of the cortical surface through the transparent skull preparation in an example Thy1-GCaMP6s mouse. M1 injury site indicated by red asterisk; black dashed line indicates border of cement barrier used to reduce spread of bleeding. The D2 ROI encompasses the mapped location of the D2 whisker within S1 barrel cortex. Black rectangle indicates a ROI used for comparison in subsequent panels. **B,** Calcium signal time series from ROIs in A. Dotted lines in time series data indicate periods of interpolated data between sequentially recorded movies. Arrows indicate time of M1-CCI injury. Background colors indicate the signal phases (Baseline, Initial Response [IR], Wave, Depression, Recovery; see legend at left) identified for further analysis. **C,** Immediate effect of M1-CCI injury on widefield cortical calcium signals. Single frames at time points indicated at top of frame show the temporal progression of GCaMP6s fluorescence after M1-CCI. Top row, Raw grayscale fluorescence (arbitrary units); Bottom row, relative change in fluorescence (% ΔF/F; pseudocolored). Signal phase (IR, Wave, Depression, Recovery) indicated next to the time stamp. Note that insets in Depression and Recovery frames use alternate color scale. **D,** M1-CCI induced calcium signals from the D2 barrel cortex. Data overlaid for n=6 mice. **E,** ROI grid used for amplitude analysis overlaid on fluorescence image of cortical surface. **F,** Box and whisker plots of maximum calcium signal amplitudes for IR, Wave, Depression (Depr), and Recovery (Rec) phases, measured from the D2 ROI or the maximum of all ROIs (n=9 mice for D2 ROI; n=10 mice for whole frame ROI).

To characterize the sequence of CCI-induced events, we measured the time-series amplitudes of IR, Wave, Depression and Recovery in both the mapped D2-whisker related column of S1 barrel cortex, and from up to 100 evenly distributed regions of interest (ROIs) spanning the field of view. Time-series calcium signals from the D2 barrel column are shown for 6 mice in **figure 2D**, and an example ROI grid covering the cortical surface in **figure 2E**. Average values of the phases of the CCI-induced cortical response are shown in **figure 2F**. Note that the peak ΔF/F values of up to 350% were 50-100 fold larger than spontaneous or sensory-evoked activity, indicating a pathophysiological response to CCI (further quantified on the cellular level in figure 5).

These results demonstrate that the cortex responds to a localized CCI in M1 with a stereotyped sequence of events, including an initial rapid calcium signal at the injury site, followed by a massive calcium signal Wave that spreads slowly across the dorsal cortex. After the Wave, the signal shows Depression with variable tendency for Recovery.

### Propagation of widespread cortical calcium signals after focal injury: spatiotemporal characteristics

To further characterize the spatial properties of the global cortical response to focal CCI of M1, we measured a number of additional parameters, including the spread directions from the injury site, the amplitude dependence on distance, maximum amplitude trajectories, and relative localization. Analysis was restricted to the cortical hemisphere ipsilateral to the CCI site because we found signals related to the IR, but no signals related to the Wave, Depression or Recovery, in the contralateral hemisphere.

We chose ROIs along four directions from the M1 injury site: anterior-posterior 0°, anterior-posterior and medial-lateral 22.5°, 45° and 67.5° (**Fig. 3A**). As shown in example data, the amplitudes of IR, Wave, and Depression varied with increasing distance along a given direction from the injury site (**Fig. 3B, C**). Notably, in this example, the Wave increased in amplitude with greater distance from the injury site. Signal amplitudes tended to be larger for the 45° and 67.5° angle directions, corresponding to lateral rather than posterior cortical areas (**Fig. 3D**). To analyze calcium signal amplitudes as a function of distance from the injury site, we averaged data for all four directions in each mouse (**Fig. 3E**). IR and Depression showed significant decreases in normalized amplitude with distance, indicating that these signals were largest near the injury site. Remarkably, wave amplitude showed no change with distance, indicating that the slow wave maintains its amplitude for up to 5.7 mm across cortex. Recovery amplitude also did not change with distance. ANOVA of LME model of normalized amplitude over distance from the ROI 2.2 mm from injury site for Initial Response (F=14.83, p=9.81e-08), Wave (F=1.06, p=0.41), Depression (F=10.61, p=1.82e-05), and Recovery (F=0.711, p=0.621); Holm corrected multiple comparisons showed significant differences between ROI’s, * – 2.2 mm, ^ – 2.9 mm, x – 3.6 mm, from injury site,* p <0.05, ** p<0.01, *** p< 0.001.

**Figure 3.**
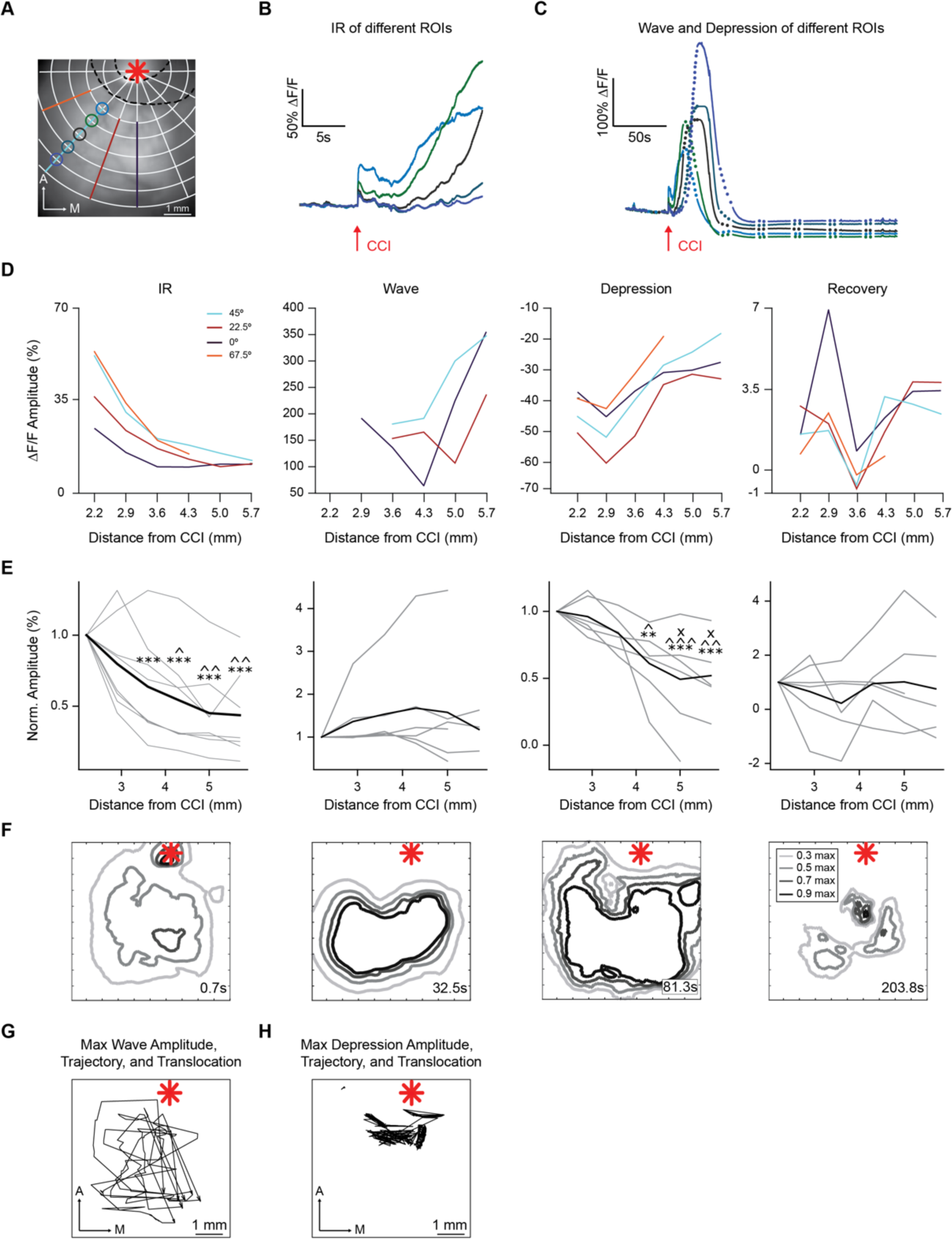
Spatial organization of cortical calcium signals induced by M1-CCI. **A,** Positions of ROIs for directional analysis (four directions: 0, 22.5, 45, 67.5 degrees from midline) overlaid on fluorescence image of cortex. M1 injury site indicated by red asterisk; black dashed line indicates border of cement barrier used to reduce spread of bleeding. **B,** Example calcium signals from five ROIs along the 45 degree direction in A. Arrow indicates time of M1-CCI. The compressed time scale highlights the IR and Wave onset. **C,** Same data in B on an expanded time scale to illustrate Wave and Depression following the IR. Dots represent periods of interpolation between sequentially recorded movies. **D,** Calcium signal amplitude (% ΔF/F) as a function of distance along four directions from M1-CCI injury site. Directions indicated by color, as shown in legend. Data from same example mouse as above. **E,** Normalized calcium signal amplitude as a function of distance from M1-CCI injury site. Thin lines represent individual mouse data averaged from all four directions; thick lines represents group means (n=8-6, 8 for IR, and 6 for W, D, and R). ROI’s, * – 2.2 mm, ^ – 2.9 mm, x –3.6 mm, from injury site, * p <0.05, ** p<0.01, *** p< 0.001. **F,** Contour plots of the IR, Wave, Depression, and Recovery signals evoked by M1-CCI. Example data from the same subject as A-D. Time post-CCI indicated above each plot. Contours correspond to the spatial profile of signals at 0.3, 0.5, 0.7, and 0.9 maximum. M1 injury site indicated by red asterisk. **G,** Direction of Wave propagation. Lines with arrows show trajectories of calcium signal maximum amplitude (ΔF/F; thin lines) for individual subjects (n = 7 mice). Position of maximum amplitude measured once per second, with interpolated periods shown as lines. M1 injury site indicated by red asterisk. **H,** Similar analysis as in G, but for trajectories of peak ΔF/F reduction for Depression (n = 7 mice).

Contour plots at near maximal time points for each signal type revealed additional spatial features (**Fig. 3F**). At 0.7 s after impact, the IR signal was localized near the injury site but also showed diffuse activation sites, possibly from the response to mechanical deformation of the cortical surface. Wave and Depression contours indicated a large coverage of the cortical surface at their respective peaks of 32.5 and 81.3 s post-injury. By contrast, Recovery signals appeared spatially less uniform, in addition to their higher variability of occurrence (as mentioned above). We determined the direction of signal spread in each mouse by tracking the position of peak signal amplitude at 1 s (100 frame) intervals. The Wave propagated mainly in the posterior-medial direction over the entire extent of the imaged cortex (**Fig. 3G**), whereas the Depression (negative peak signal) remained localized to areas closer to the injury site (**Fig. 3H**). To compare the relative localization of the signals, we measured the distances between the geometrical center coordinates of activation in individual animals (80% maximum amplitude threshold over total imaged area for IR, Wave and Depression; 10% maximal Depression amplitude threshold for Recovery). All distances between the center positions of IR, Wave, Depression and Recovery were significantly different from 0, indicating spatially segregated cortical locations of the different phases of peak CCI-evoked signals. Peak signal locations were mainly within somatosensory, visual, and association cortical areas (see also figure 6 for further relation to cortical areas). These data show that the phases of cortical signals following focal injury to M1 have distinct spatiotemporal features. The IR decayed quickly, followed by a Wave that propagated without amplitude decrement over the entire extent of the imaged cortex. The signal later showed Depression and, in some cases, Recovery. The amplitudes of IR, Wave, Depression, and Recovery signals peaked at distinct cortical locations.

To visualize the timing of the CCI-evoked signals, we constructed activation time maps (**Fig. 4A**), where color reflects the time of calcium signal amplitude reaching a threshold (50% peak amplitude for IR and Wave, 80% minimum amplitude for Depression, and 10% difference from Depression for the Recovery). Note that the color scales are compressed or stretched to match the time scale of the signals. For example, the IR signal is shown from 0 to 0.3 s, while the Wave is shown from 0 to 84 s. Signals spread in a posterior direction from the M1 injury site, with distinct but overlapping spatial profiles. ROI analysis of group data (**Fig. 4B**) confirmed the delayed temporal progression of signals from the injury site to posterior cortical areas. Analysis of the activation areas was done by masking pixels that exceeded a threshold of 20% of peak amplitude for the IR and Wave, and Depression, and 10% of Recovery (**Fig. 4C**). The time course of area changes for individual mice showed dynamic increases and decreases of the activation areas for each signal type (**Fig. 4D**). IR returned to baseline within 1s, while activation area of the Wave lasted up to 60 s, and Depression lasted even longer. The average peak area changes are shown in **figure 4E**.

**Figure 4.**
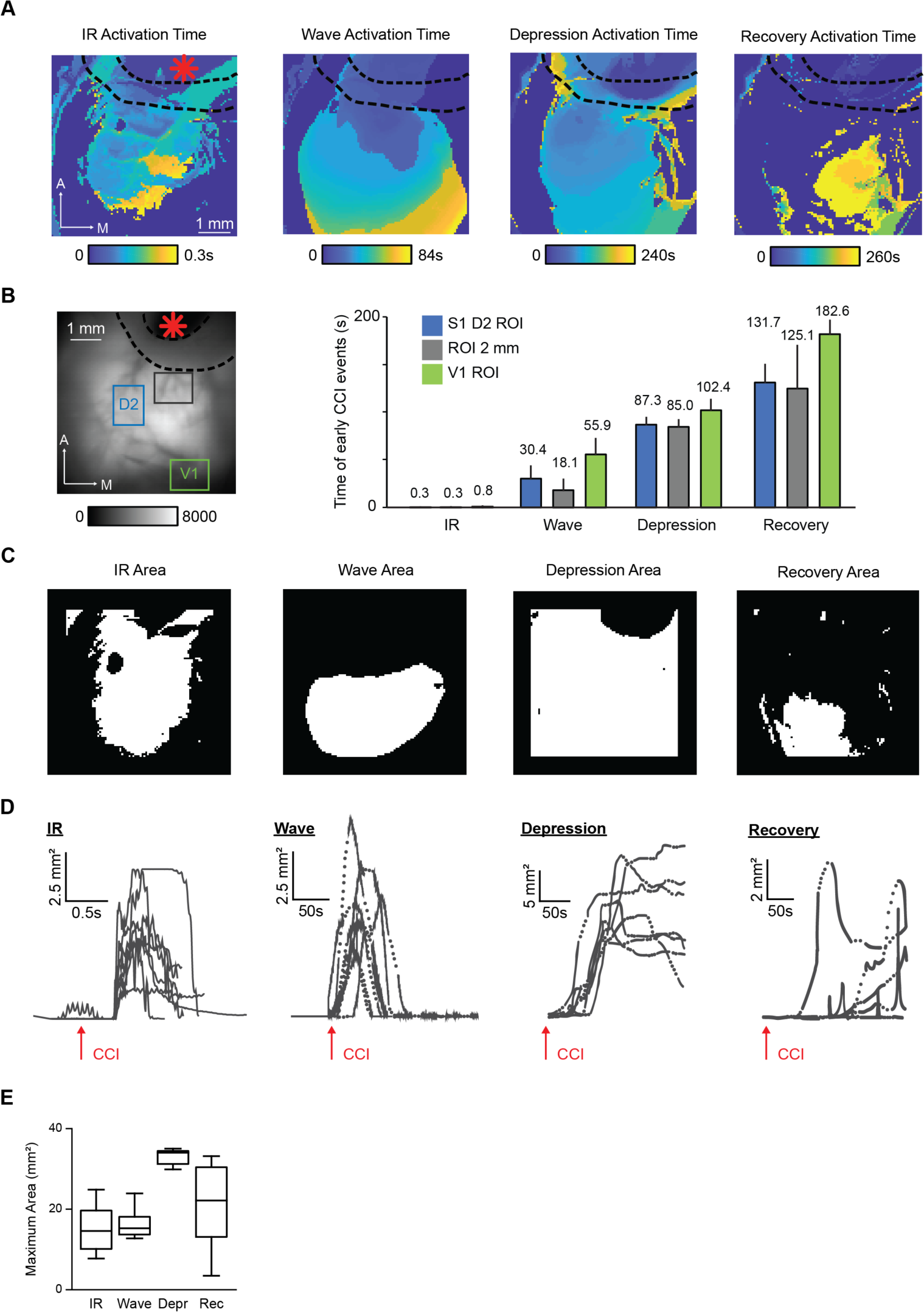
Temporal and spatial organization of cortical calcium signals induced by M1-CCI. Calcium signal activation time, area and propagation. **A,** Example activation time maps. Color represents the time of calcium signal threshold crossing (50% peak amplitude for IR and Wave; 80% of minimum amplitude for Depression; 10% difference from Depression for the Recovery. Red asterisk indicates M1 injury site; black dashed line indicates cement barrier to prevent bleeding. **B,** Left, ROIs corresponding to D2 region of S1 barrel cortex (blue), visual cortex (green), a site 2 mm from M1 injury (brown), overlaid on fluorescence image of cortical surface through transparent skull preparation. Right, Plot of the mean time to peak for each type of signal in the three selected ROIs. Data are mean ± SD (N mice by column = 7,7,8,7,7,8,7,8,7,4,3,2). Recovery has lower N because in some cases Recovery onset was not captured by video recording or did not occur on the time scale of our measures. **C,** Example area maps for the IR, Wave, Depression, and Recovery phases of M1-CCI evoked signals. Calculated based on the active pixels at 20% peak threshold (for IR, Wave and Depression) and 10% of recovery. **D,** Time series of area dynamics for individual mice (n = 7) for IR, Wave, depression and recovery, individual traces. Time of injury marked with arrow. Dotted lines in time series data indicate periods of interpolated data between sequentially recorded movies. **E,** Box and whisker plots of maximal area of M1-CCI evoked signals (n = 10 mice for IR; n = 7 mice each for Wave, Depression, Recovery).

### Two-photon calcium imaging of S1 neuronal population activity during focal injury of M1

To investigate the effects of cortical injury with cellular resolution, we performed two-photon calcium imaging in S1 during M1 CCI (**Fig. 5A, B**). Neuronal populations in layer 2/3 of the D2 barrel column were imaged through a chronic cranial window implanted above the S1 barrel field, and CCI was applied to M1 in awake, head-fixed Thy1-GCaMP6s mice (as above for the widefield experiments). Spontaneous activity and D2 whisker-evoked sensory activity were recorded during four baseline sessions before the CCI session. An example response of S1 neurons to M1 CCI is shown in **figure 5C and D**. In the time series fluorescence changes, we observed a massive calcium signal increase with a delayed onset after the CCI, corresponding to the Wave (**Fig. 5C**). Signals related to the IR were not detected. Data from ROIs corresponding to the whole imaging frame or to individual neurons showed consistent fluorescence increases during the Wave, suggesting that both neurons and the surrounding neuropil underwent strong activation. However, for individual neurons, we observed variability in the duration of Wave-related activation before the calcium signals returned toward baseline, with some neurons showing elevated signals for tens of seconds longer than other neurons (**Fig. 5C**). Framewise images of the Wave propagation through the two-photon field of view revealed key aspects of cellular activation (**Fig. 5D**). First, neuronal somata showed activation at the leading edge of the Wave before being saturated by the strong fluorescence of the Wave. The wavefront of the calcium signal spread in a continuous fashion (**Fig. 5D, top row**), consistent with the widefield imaging data (above). Both “silent” neurons (neurons that were not observed to be active in previous sessions) and active neurons became activated during the Wave. During the onset of the Depression, neurons gradually showed reduced activity, with some neurons remaining more persistently active (**Fig. 5D, bottom row**). During the later phase of the Depression, nearly no neurons were active (**Fig. 5E, left**). Finally, we observed in a small number of neurons a “reactivation” of the large calcium signal Wave during the later phase of the Depression (**Fig. 5E**). The large amplitude of this reactivation signal was more similar to the Wave than to spontaneous or sensory-evoked activity, and could be either short- or long-duration (**Fig. 5E, F**). The effects of M1 CCI on S1 neuronal population activity are summarized in **figure 5F**. Note that the Wave recruited a much larger percentage of active neurons compared to spontaneous and whisker-evoked activity and that the peak signal was an order of magnitude larger in amplitude (**Fig. 5F**), highlighting the dominant effect of the Wave on S1 population activity.

**Figure 5.**
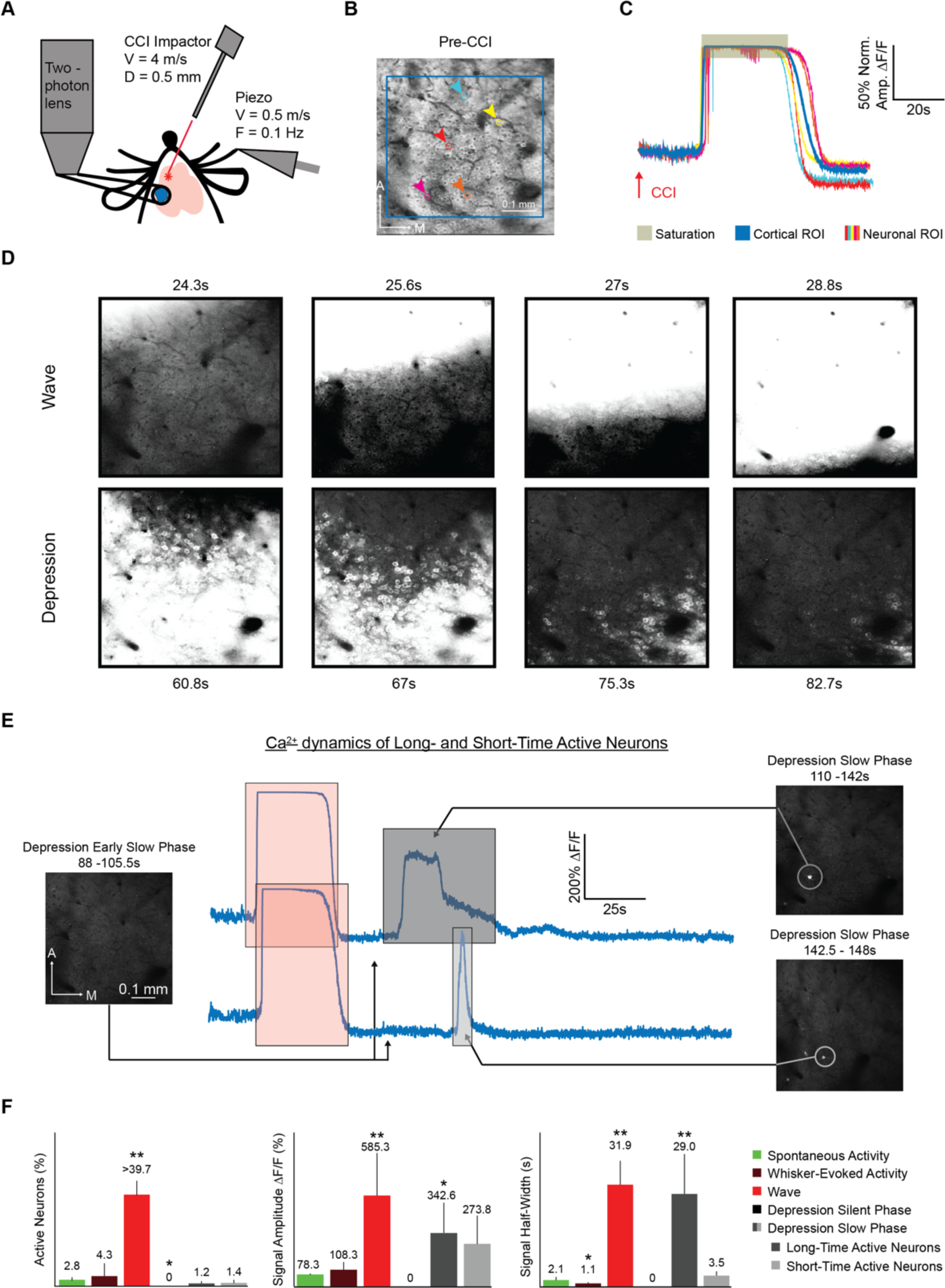
Two-photon calcium imaging of cortical activation during M1-CCI focal injury. **A,** Schematic of experimental setup. Velocity (V) and diameter (D) of CCI probe are the same as in earlier figures. V and frequency (F) are shown for the piezoelectric actuator (Piezo) used for whisker deflection. Blue circle indicates imaging field of view containing the D2 whisker area of S1 barrel cortex. Red asterisk represents M1 injury site. **B,** Two-photon fluorescence image of S1 L2/3 neuronal populations through a chronically implanted glass window in a Thy1-GCaMP6s reporter mouse (average of 100 frames, 15.49 frames/s). Selected individual neurons are marked with colored circles. Blue rectangle marks “Cortical ROI” for larger region comparison with cellular signals. **C,** Example calcium signals from ROIs in B. Cellular and larger area ROIs shown by thin and thick lines, respectively). Signals are normalized to peak in order to compare timing. Arrow indicates time of M1-CCI. **D,** Temporal progression of S1 cellular and neuropil fluorescence changes after M1-CCI. Time post-CCI indicated above each image. Each image represents the average of 15 consecutive frames (15.46 frames/sec; grayscale, arbitrary units). Field of view is the same as in B. **E,** Individual neuron activity in the slow phase of post-Wave Depression. Example calcium signals of two neurons during the Depression phase that follows the Wave. While overall fluorescence was below baseline, two neurons showed a secondary reactivation of large-amplitude, long-duration signals. Same field of view as above. **F,** Summary of cellular calcium signals induced by M1-CCI. Mean changes in percent of active neurons (left), peak amplitude (middle), signal duration (width at half height; right). Spontaneous activity and whisker-evoked signals are also included for comparison (color code shown in legend). Values are mean ± SD for n>=10 cells from n>=4 mice in each condition. Mean values are listed above bars. Significance indicated by: * p <0.05, ** p<0.01; measured with paired t-test).

The neuronal wave amplitude and duration were 586±270 ΔF/F% and 31.8±7.3 s full-width at half-maximum (FWHM) (mean ± SD, n=4 mice). For spontaneous activity, signal amplitude and duration were 78.3±8.6 ΔF/F% and 2.1±0.8 s FWHM. For sensory-evoked activity, amplitude and duration were 108.3±44.1 ΔF/F% and 1.1±0.2 s (mean ± SD, n=8 mice) respectively. The percent of active neurons during the Wave was increased from less than 5% during spontaneous (2.8±1.2% and 4.3±5.3% for spontaneous and whisker-deflection evoked activity respectively; mean ± SD, n=8 mice) to at least 39.7±6.0% (mean ± SD, n=4 mice). During the slow phase of Depression the “unusual” activity of individual neurons activity was 343±207% in amplitude and 29.0±12.2 s FWHM in duration for long-time active cells, and 274±177% and 3.5 ±1.1 FWHM for the short-time active cells. The percentage of active neurons during the slow phase of Depression was 1.2±0.7% for spontaneous and 1.4±1.1 for whisker-evoked activity, respectively (mean ± SD, n=4 mice).

To summarize results thus far, widefield calcium imaging showed a large-amplitude, global, and long-lasting cortical calcium signal evoked by CCI applied to the ipsilateral M1. The consistent events that followed M1 CCI were a fast IR, a slower and more spatially extensive Wave of maintained amplitude, followed by an equally spatially extensive Depression, and a variable Recovery of the calcium signal to pre-injury baseline. While the overt features of the CCI-induced cortical signals were consistent across mice, the within-subject signal parameters were generally uncorrelated (for example, IR amplitude only predicted signal Recovery, not Wave or Depression amplitude), suggesting a common pattern of cortical activation in response to focal injury with subtle individual differences in signal propagation. Two-photon calcium imaging experiments confirmed the basic features of the CCI-evoked signals while providing further insight into the cellular basis of these signals. A large fraction of S1 cortical neurons were activated during the Wave, with subsets of individual neurons showing longer lasting Wave-related activity or post-Wave reactivation.

### Longitudinal measurements of sensory maps, cortical network connectivity, and behavior after M1-CCI

We aimed to follow the effects of local M1 injury on cortical function and behavior over the subsequent days and weeks. To determine the lasting effects of M1-CCI on cortical function, we tracked spontaneous and sensory-evoked cortical activity on days 1, 3, 7, 14 and 56 post-CCI. Baseline measures were acquired in four pre-CCI sessions (days −7, −6 and −4; as in timeline of figure 1). In each session, whisker deflections were delivered to the contralateral D2 whisker of awake, head-fixed Thy1-GCaMP6s mice (**Fig. 6A**), and trial-averaged cortical functional maps acquired with widefield calcium imaging (**Fig. 6B**). Time series data from a ROI encompassing the D2 barrel column (**Fig. 6C**), as well as spatial parameters from the cortical maps were used for analysis of signal parameters. An example of repeated imaging of whisker-evoked cortical maps for an example subject is shown in **figure 6D**, where changes in the spatial profile are evident at early time points following M1-CCI.

**Figure 6.**
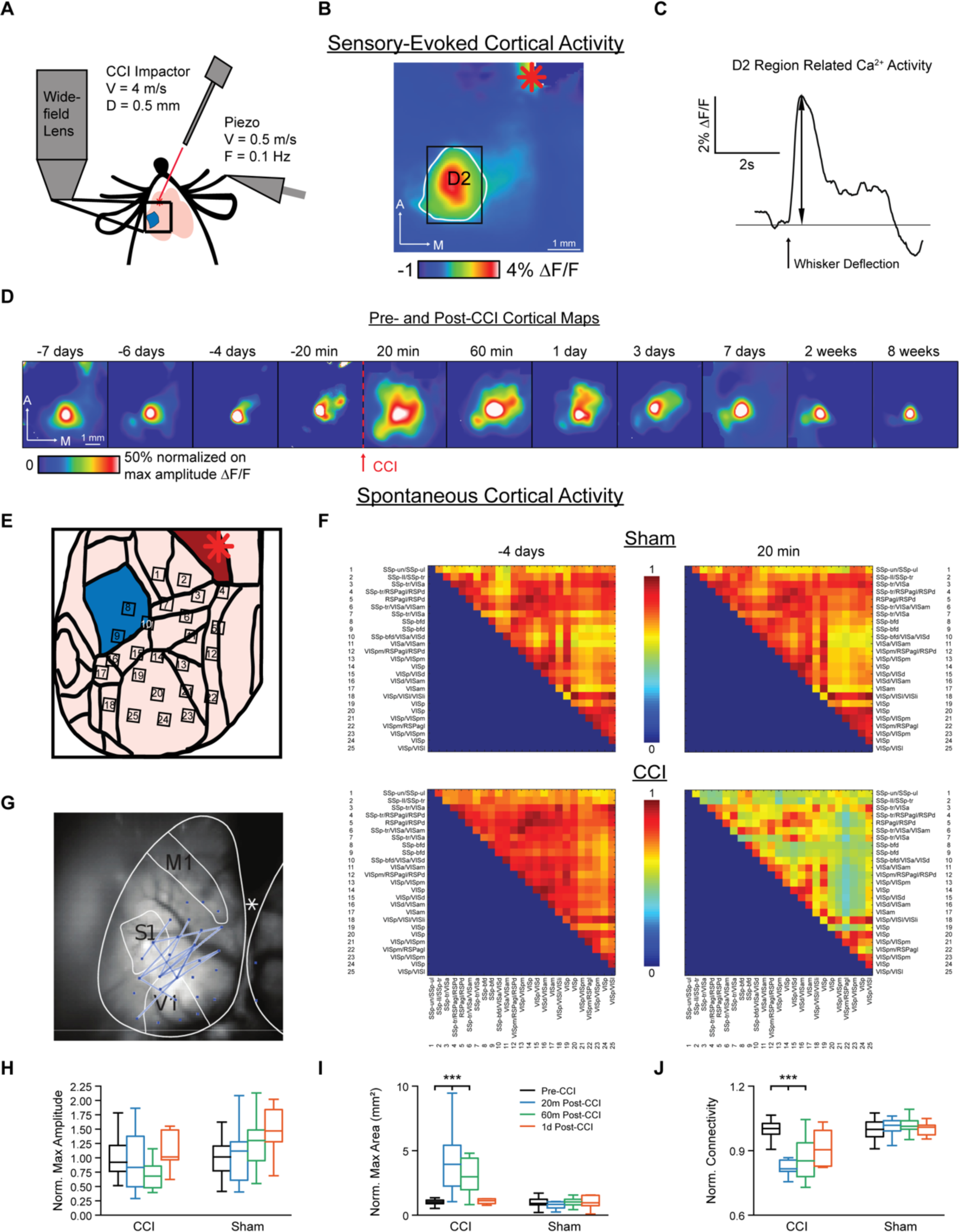
Effects of M1-CCI on sensory maps and cortical functional connectivity. **A,** Schematic of experimental setup for widefield imaging of sensory-evoked and spontaneous cortical calcium signals in Thy1-GCaMP6s mice. CCI and sensory stimulation parameters are the same as in figure 5. **B,** Example sensory map in response to D2-whisker deflection under pre-CCI baseline conditions. Calcium signal fluorescence (ΔF/F) averaged for 30 trials with warmer colors representing larger relative changes. White outline indicates area of signal; black rectangle indicates ROI used for time series extraction. Red asterisk shows M1 injury site. **C,** Example time series of sensory-evoked calcium signal, corresponding to data in B. Time of D2 whisker deflection marked with thick black arrow. Signal amplitude marked with thin black arrow. **D,** Repeated measurement of sensory maps before and after CCI from one subject. Time points relative to CCI shown above each frame. Maps are normalized and scaled to 50% maximal amplitude for comparison across time points. **E,** ROIs used for functional connectivity measures, shown superimposed on schematic cortical areas based on Allen Mouse Brain Common Coordinate Framework (https://scalablebrainatlas.incf.org/mouse/ABA_v3). Red asterisk indicates M1 injury site. **F,** Group-averaged functional connectivity matrices measured from spontaneous cortical calcium signals, 4 d before (left) and 20 min after (right) M1-CCI. Sham group, top row; Injury group, bottom row. Spontaneous activity measured over 160 s (8 movies, each 20 s duration) of Injury (n=12 mice; 7 male, 5 female) and Sham groups (n=11 mice; 7 male, 4 female). **G,** Changes in functional connectivity between −4d and +20 min time points mapped onto cortical ROIs. Blue lines connect ROIs with changes common across mice in the Injury group. **H,** Changes in sensory-evoked calcium signal amplitude after M1-CCI for Injury and Sham groups. Amplitude data (n = 12 and 11 mice for injury and sham groups respectively) are shown for pre-CCI, and 20 m, 60m, 1d post-CCI. **I,** Changes in sensory-evoked calcium signal area after M1-CCI for Injury and Sham groups, shown for 20m pre-CCI, and 20m, 60m, and 1d post-CCI. **J,** Changes in normalized functional connectivity measured from spontaneous activity after M1-CCI for Injury and Sham groups, shown for baseline, 20m, 60m, and 1d post-CCI, as in panels H and I.

We measured spontaneous widefield cortical activity in the same imaging sessions, and applied network analysis methods to determine changes in functional connectivity between cortical areas after M1-CCI (Salsabilian et al., 2022). We used the group-averaged connectivity to determine cortical network changes at the global level. Adjacency matrices were calculated from time series data for each trial using the ROIs shown in **figure 6E**, and ROIs were overlaid onto the Allen brain atlas to generalize the cortical areas (Allen Mouse Brain Common Coordinate Framework, https://scalablebrainatlas.incf.org/mouse/ABA_v3). For each recording session, the connectivity network in both the Injury and the Sham groups were computed by averaging the adjacency matrices across subjects and trials within each group. As an example, **figure 6F** shows the group-averaged connectivity networks for one pre- and one post-CCI (or Sham) session for each group. Note that the baseline (−4 days) group-averaged network in both the Injury and the Sham groups show similar connectivity, especially between SS and VIS areas (**Fig. 6F**). While the Sham group showed no obvious change between the time points (**Fig. 6F, top row**), most connections (or links) between ROIs in the Injury group showed decreased mean connectivity (**Fig. 6F, bottom row**). To determine which ROI pairs significantly contributed to the altered connectivity networks before and after focal injury, the student t-test was applied to the connectivity networks of pre- and post-CCI sessions. Links with significant differences common across subjects are shown for the −4 days pre-CCI and 20 min post-CCI sessions in **figure 6G**. The links correspond mostly to connections between the somatosensory (SS) and visual (VIS) regions identified from the Allen brain atlas. The widespread changes in connectivity were consistent with the localizations of the peak IR, Wave, Depression and Recovery signals (Peak IR signals localized to SSp-II, SSp-tr, VISa; peak Wave signals, VISam,pm,p and RSPagl; peak Depression signals, SSp-tr, VISa,d,p and RSPagl; peak Recovery signals, RSPagl, VISpm,p).

The group mean effects of M1-CCI on sensory-evoked cortical maps and network connectivity (spontaneous activity) are shown in **figure 6H, I, J**. The peak amplitude of D2 whisker-evoked cortical maps did not significantly increase in the days following CCI, as did occur in the Sham group (**Fig. 6H**). ANOVA of LME model (Amplitude ∼ Condition * Time + Sex + (1|Mouse)) (F=3.68, p=0.068). However, large changes in the area of sensory-evoked cortical maps occurred after CCI that were not apparent in the Sham group (**Fig. 6I**). ANOVA of LME model (Area ∼ Condition * Time + (1|Mouse)) showed a significant effect of the Condition:Time interaction (F=6.89, p=1.82e-08). Post hoc Holm corrected MCT of Condition and Time interaction of sensory evoked area of calcium activity between Injured and Sham groups 20m (t=6.45, p=2.03e-07) and 60m (t=4.77, p=0.00063) post-CCI showed significant difference at early time points (before 1d). The normalized area of sensory-evoked maps increased by 4.40-fold at 20 minutes post-CCI and remained at 2.58-fold greater than baseline in combined post-CCI time points (20 min, 60 min, 1 d, 3 d, 7 d, 14 d). We also found changes in group-averaged CCI-induced cortical network connectivity, measured from spontaneous activity. While the Sham group showed no significant change in mean connectivity, the Injury group showed a significant decrease in mean connectivity at both the 20 minute (0.83 normalized to pre-CCI) and combined post-CCI time points (0.93 normalized to pre-CCI) (**Fig. 6J**). ANOVA of LME model (Connectivity ∼ Condition * Time + (1|Mouse)) showed a significant effect on the Condition:Time interaction (F=13.36, p<2e-16). Post hoc Holm corrected MCT of Condition and Time interaction of spontaneous connectivity of calcium activity between Injured and Sham groups 20 min (t=-6.94, p=1.91e-08) and 60 min (t=-5.28, p=7.43e-05) post-CCI showed significant difference at early time points (before 1d). Together these results indicate that M1 CCI leads to sensory maps of increased area without a change in amplitude, and also leads to decreased functional connectivity between cortical regions. Effects were strongest at early post-injury time points and tended to recover over subsequent days, as analyzed further below.

### Time course of behavioral, activity and connectivity changes after focal injury of M1

**Figure 7** shows Sham and Injury group mean time course data for behavior, sensory-evoked maps, and cortical connectivity for four baseline pre-CCI (−7 d, −6 d, −4 d, −20 min) and 7 post-CCI (20 min, 60 min, 1 d, 3 d, 7 d, 14 d, 56 d) time points. For the behavioral measures, observations of grooming and rearing incidents were made for each mouse during the 10 min after each wide-field imaging session when the mouse was returned to the home cage. After M1-CCI, we observed strong immediate reductions of both rearing and grooming in the Injury group compared to Sham, although the Sham group also showed smaller post-Sham reductions in rearing and grooming (**Fig. 7A**). Comparison of the Sham and Injury groups showed significant differences in rearing and grooming at the 10 min time point. ANOVA of LME model (∼ Condition * Time + (1|Mouse)) showed significant interaction with injury and time for rearing (F=3.51, p=0.00015) and grooming (F=2.279, p=0.012), n=12 mice (7 male, 5 female) in injury group and 11 mice (7 male, 4 female) in Sham; post-hoc Holm corrected MCT showed significant difference between injured and sham for rearing (t = −4.096, p = 0.011) and grooming (t=-3.95, p=0.028) at 10m post-CCI. Rearing and grooming recovered to pre-CCI levels within 1 d after injury. Separate analysis of male and female mice revealed no sex-specific behavioral effects of M1-CCI, F=(0.976, 0.077), p=(0.335, 0.784) for rearing and grooming respectively.

**Figure 7.**
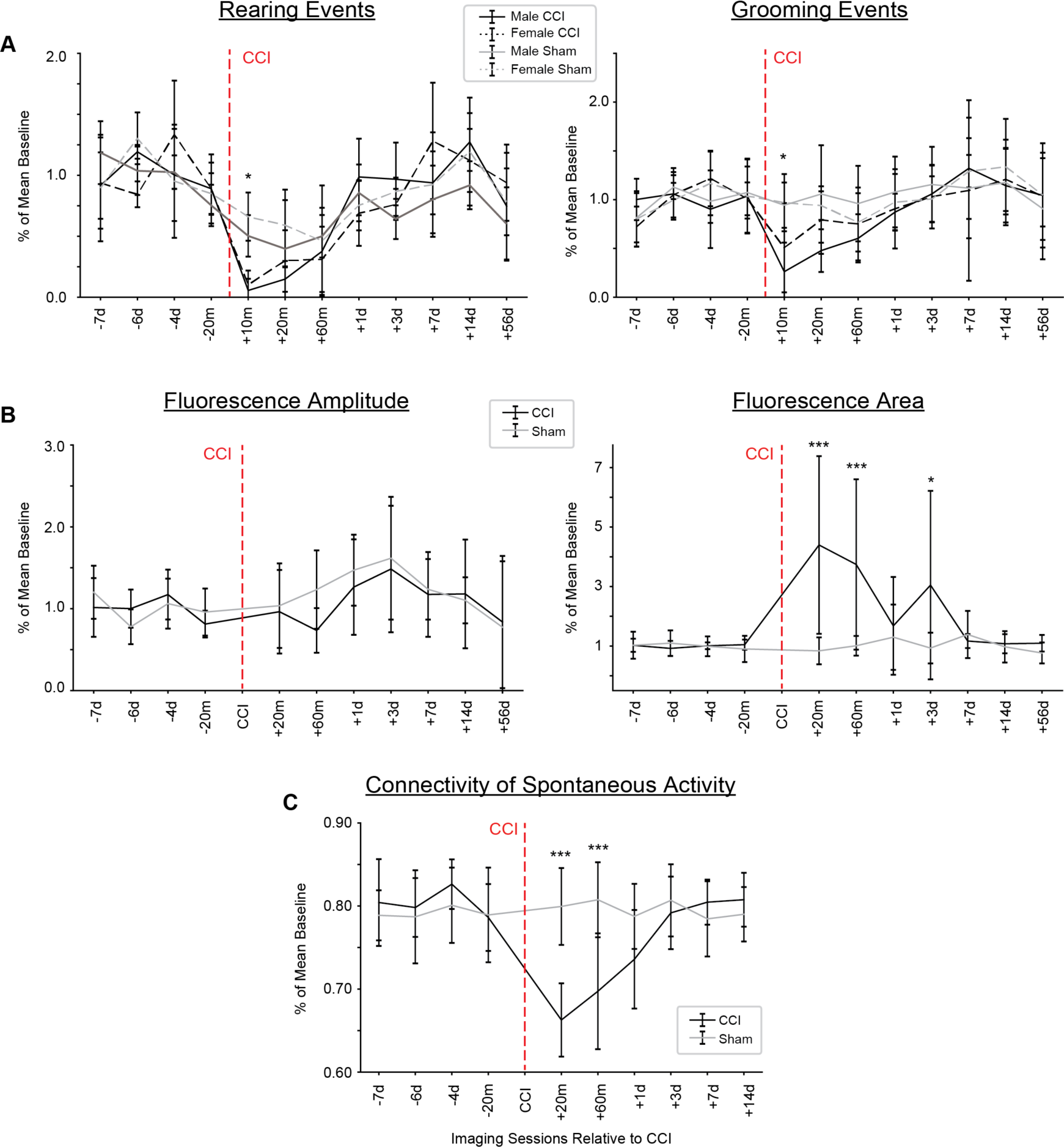
Time course of changes in sensory maps, functional connectivity, and behavior after M1-CCI. **A,** Effect of M1-CCI on rearing (left) and grooming (right) behavior. Observations were made in the 10 min after imaging sessions when mice were returned to home cage. Mean (± SD) number of observations for each group normalized to pre-CCI baseline mean. Black line, Injury group; Gray line, Sham group. Dotted lines indicate female mice. Time of injury marked with an arrow, * p < 0.05. **B,** Effect of M1-CCI on sensory-evoked calcium signal amplitude (left) and area (right), from widefield imaging in Thy1-GCaMP6s mice. Mean (± SD) values for each group normalized to pre-CCI baseline mean. Black line, Injury group; Gray line, Sham group. Time of injury marked with an arrow. Changes in sensory-evoked calcium signal amplitude and area after M1-CCI for Injury and Sham groups. Amplitude data (mean ± SD; n =12 and 11 mice for injury and sham groups respectively) are shown for 20m pre-CCI, and 20 m, 60m, 1d, 3d, 7d, 14d post-CCI. Changes in sensory-evoked calcium signal area after M1-CCI for Injury and Sham groups. Area data (mean ± SD) are shown for pre-CCI, and 20m, 60m, 1d, 3d, 7d, 14d post-CCI. **C**, Effect of M1-CCI on functional connectivity during spontaneous activity from widefield imaging in Thy1-GCaMP6s mice. Mean (± SD) values for each group normalized to pre-CCI baseline mean. Black line, Injury group; Gray line, Sham group. Time of injury marked with an arrow. In all panels, days with significant changes compared to baseline are marked with asterisk (* p <0.05, ** p<0.01, *** p<0.001).

Time course analysis of sensory-evoked widefield cortical maps (**Fig. 7B, left**) showed a trend of increased signals in both Injury and Sham groups. There was, however, no significant difference between the amplitudes of sensory-evoked widefield cortical maps in Injury and Sham groups (ANOVA of LME model (∼ Condition * Time + (1|Mouse)) (F=0.979, p=0.462)). For cortical map area measures (**Fig. 7B, right**), we found large increases in the Injury but not Sham condition. Map area was significantly larger for mice in the Injury group at the 20 min, 60 min, and 3 d post-CCI time points (ANOVA of LME model (∼ Condition * Time + (1|Mouse)) showed significant interaction with injury and time for normalized calcium area, F= 6.76, p = 5.41e-09, n=13 mice in injury group and 11 in Sham; adjusted Holm test showed significant differences between injured and sham 20 min, 60 min and 3 days after injury, p = 4.09e-08, 0.00024, 0.0395 respectively). Similar to behavioral data, sex-specific analysis of sensory-evoked map area showed no significant difference between males and females (ANOVA of normalized area of sensory-evoked activity to sex; F = 1.357, P = 0.256).

Time course analysis of mean network connectivity (**Fig. 7C**) showed significantly decreased connectivity at the 20 m and 60 m post-CCI time points (ANOVA of LME model (∼ Condition * Time + (1|Mouse)) showed significant interaction with injury and time for normalized connectivity, F =12.75, p =9.81e-16, n=13 mice in injury group and 12 in Sham; adjusted Holm test showed differences between Injured and Sham groups 20 m and 60m after injury, p = 4.84e-08 and 0.00023 respectively). Mean connectivity recovered to near-baseline values by 3 d after the injury. Similar to behavioral data, sex-specific analysis of sensory-evoked map connectivity showed no significant difference between males and females (ANOVA of normalized area of connectivity to sex; F = 0.528, P = 0.476).

Together these data show concomitant changes in behavior, cortical sensory maps, and network connectivity in the minutes, hours, and days following focal injury to M1.

### Repeated two-photon imaging of S1 neuronal population activity after focal injury of M1

We found earlier in the study (see Fig. 5), using two-photon calcium imaging, that the number of active neurons increased during the Wave and decreased during the Depression. How does M1-CCI affect whisker-evoked activation of neuronal populations in layer 2/3 of S1? To address this question we performed chronic two-photon calcium imaging in awake, head-fixed Thy1-GCaMP6s mice at multiple time points before and after focal injury (**Fig. 8A**). Mechanical stimulation of the D2 whisker was delivered with a piezoelectric wafer in the same way as in the widefield imaging experiments. Example images from the same field of view 20 minutes before, 20 minutes after, and 14 days after M1 CCI are shown in (**Fig. 8B)**. Cellular ROIs were identified (**Fig. 8C**) and time series data extracted and compared for each imaging session. Qualitatively, the sensory evoked and spontaneous cellular calcium signals appeared similar before and after M1-CCI (**Fig. 8D**). Indeed, analysis of group data indicated that the largest effect of CCI was a reduction of the number of active neurons for both sensory evoked and spontaneous activity (**Fig. 8E**). The reduction in the number of active neurons was strongest at the shortest post-CCI time point (20 min) and remained significantly reduced in pooled post-CCI time points (20 min, 60 min, 1 d, 3 d, 7 d, 14 d) (ANOVA of LME model (∼ Experiment * Injury + (1|Mouse)) (F=39.98, P=3.85e-09) adjusted Holm test showed differences between pre and post CCI groups of active cells (t=-6.249, p = 5.54e-09). We did not observe significant changes in the amplitude and duration (width) of the calcium signals, but we did see significant increases in event frequency (**Fig. 8F**), (ANOVA of LME model (∼ Experiment * Injury + (1|Mouse)) (F=135.58, P=< 2.2e-16) adjusted Holm test showed differences between pre and post CCI groups of event frequency for both spontaneous (t=4.711, p = 1.48e-05) and evoked (t=19.139, p = < 2e-16) active cells. We also measured the average pairwise Spearman’s Correlation of cells within imaging sessions and saw significant interaction between injured state and experiment (**Fig. 8G**) (ANOVA of LME model (∼ Experiment * Injury + (1|Mouse)) (F=546.67, P=< 2.2e-16). We found differences between pre and post CCI groups in average correlation for both spontaneous (t=5.554, p = 1.67e-07) and sensory-evoked (t=-25.635, p = < 2e-16) (adjusted Holm test) active cells. There was a significant decrease in correlations in sensory-evoked sessions post-CCI and a significant increase in correlated activity in spontaneous sessions post-CCI.

**Figure 8.**
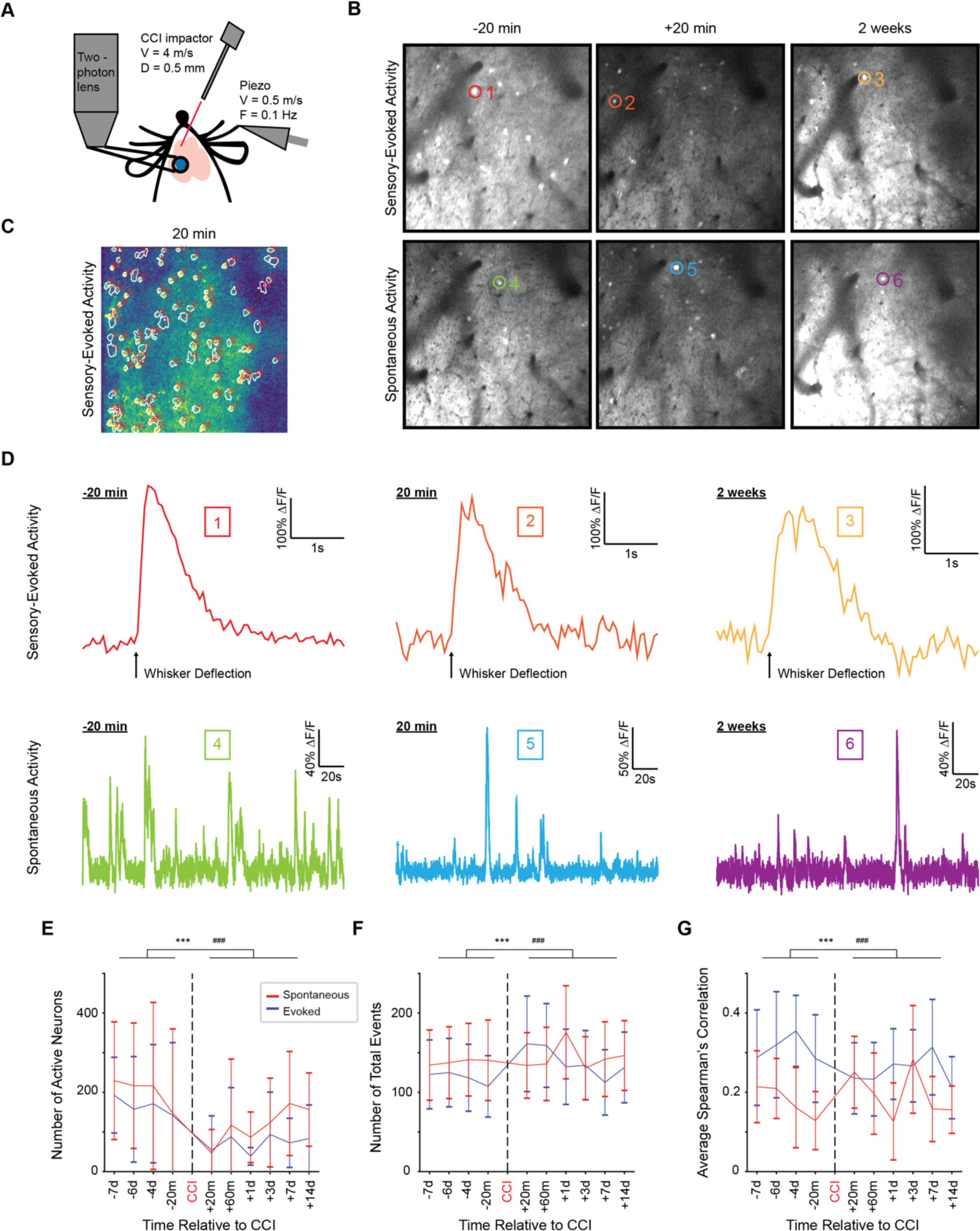
Effects of M1-CCI on S1 neuronal population activity. **A,** Schematic of experimental setup. Two-photon calcium imaging in awake, head-restrained Thy1-GCaMP6s mice before and after M1-CCI. CCI and sensory stimulation parameters are the same as in previous figures. Blue circle indicates cranial window for chronic optical access. **B,** Example two-photon fluorescence images of L2/3 neurons in the D2 barrel field of S1. Mean of 16 frames (15.49 frames/s) around periods of activity are shown for sensory-evoked (top row) and spontaneous (bottom row) activity, 20 m before, 20 m after, and 2 weeks after M1-CCI. Image processed in Image J and displayed from 0 to 30000 arbitrary units. **C,** ROIs for measuring sensory evoked activity in response to D2-whisker deflection. White outlines defined by CaImAn superimposed on a 30-trial average ΔF/F image, where warmer colors indicate larger relative change. **D,** Sensory-evoked (top row) and spontaneous (bottom row) calcium signals from example neurons, 20 m before, 20 m after, and 2 weeks after M1-CCI detected by CaImAn. Each sensory-evoked trace is the average of 33 trials; black arrow marks time of whisker deflection. Boxed numbers correspond to ROIs shown in B. It was not possible to track neurons across sessions in all mice due to clotting or field of view changes after CCI. **E,** Longitudinal mean (± SD) of active cells per mouse detected by Suite2P in sensory-evoked (Red) and spontaneous (Blue) sessions, n = 5-8 mice (n for each time point: n = 7,7,8,6,6,6,7,7,7,7 respectively) and n = 5-8 mice (n for each time point = 8,7,8,8,5,5,6,7,6,7 respectively). There were significant differences between Pre and Post CCI sessions when comparing active cells for sensory-evoked and spontaneous sessions. (Session type: * Sensory-evoked, # Spontaneous, *** p<0.001 for Pre vs Post CCI comparison). **F,** Longitudinal mean (± SD) of calcium transients of all active cells per session detected by Suite2P in sensory-evoked (Red) and spontaneous (Blue) sessions, n = 5-8 mice (n for each time point: n = 7,7,8,6,6,6,7,7,7,7 respectively) and n = 5-8 mice (n for each time point = 8,7,8,8,5,5,6,7,6,7 respectively). There were significant differences between Pre and Post CCI sessions when comparing the number of calcium transients of active cells for sensory-evoked and spontaneous sessions. (Session type: * Sensory-evoked, # Spontaneous, *** p<0.001 for Pre vs Post CCI comparison). **G,** Longitudinal mean (± SD) of mean within session Spearman’s pairwise correlation of calcium activity of active cells per session detected by Suite2P in sensory-evoked (Red) and spontaneous (Blue) sessions, n = 5-8 mice (n for each time point: n = 7,7,8,6,6,6,7,7,7,7 respectively) and n = 5-8 mice (n for each time point = 8,7,8,8,5,5,6,7,6,7 respectively). There were significant differences between Pre and Post CCI sessions when comparing within session correlation of calcium activity of active cells for sensory-evoked and spontaneous sessions. (Session type: * Sensory-evoked, # Spontaneous, *** p<0.001 for Pre vs Post CCI comparison).

These results suggest a complex cortical response to TBI on the cellular level. M1-CCI leads to a reduced number of active neurons in S1, but that when neurons are active, they display calcium signals of unchanged amplitude but increased event frequency. Furthermore, sensory-evoked cellular correlations decrease, while widefield spontaneous activity correlations increase. The mechanisms underlying these complex changes in the cellular cortical response to TBI will be an important focus of future studies.

## Discussion

We used a controlled cortical impact model of focal TBI applied to the motor cortex (M1-CCI) in transgenic GCaMP reporter mice to investigate the physiological response of the cortex to injury. We used both widefield and two-photon calcium imaging to optically record cortex-wide and cellular-resolution responses to M1-CCI, respectively. This approach allowed us to investigate the effects of focal brain injury on cortical sensory maps and resting-state functional connectivity, as well as changes in the underlying cellular activity. Our longitudinal imaging experiments uncovered injury-induced changes in spatial features of sensory maps, decreased cortical network connectivity, and reductions in the fraction of active neuronal populations in S1 sensory cortex. Most of these changes recovered, in parallel with behavioral effects, in the hours or days following injury, with the exception of a persistent decrease in the fraction of active S1 neurons.

### A note on the preparation

Our study helps to advance the experimental paradigm for investigating TBI in three basic ways. 1) Imaging in unanesthetized mice. Anesthesia profoundly affects brain state, with prominent low-frequency spontaneous activity and altered responses to sensory stimuli (Erchova et al., 2002). While technically more challenging than imaging in anesthetized mice, we acquired imaging data during application of M1-CCI in awake, head-restrained mice. This required acclimation of mice to head fixation over a period of days, and increased complexity in the experimental protocol on the day of the TBI. However, because the brain’s response to injury is likely altered by the use of anesthesia, it is important to record the response to injury under unanesthetized conditions. 2) Imaging during the induction of TBI. While previous studies have applied certain TBI protocols such as blast injury or fluid percussion in awake mice, it is difficult to acquire imaging data during induction of TBI with the mouse positioned under the microscope. Instead, alternatives using local KCl application or optogenetic approaches have been used (Monai et al., 2021; Houben et al., 2017). To allow application of CCI with head-restrained mice under the microscope, we positioned the CCI probe at an angle above the M1 injury site. This position avoided the field of view in widefield imaging experiments, and was far enough away from the microscope objective in two-photon imaging experiments. However, this experimental arrangement also introduced certain limitations. For example, the diameter of the CCI probe, and thus the extent of TBI we could produce, was limited to avoid mechanical and optical interference with image acquisition. We also limited the injury extent to avoid obscuring of the field of view from dural blood seepage and to avoid loss of consciousness during head restraint. 3) Longitudinal tracking of cortical maps and neuronal populations after TBI. While it is a valuable to understand how individual neurons and brain areas undergo long-term changes after injury, most experiments use cross-sectional designs with multiple experimental groups to establish a time course. Using chronic GECI imaging allowed us to track the progression of large-scale and cellular changes in calcium signals across spatial scales of brain function (Lee et al., 2020). While we used widefield imaging and two-photon microscopy in separate experiments, recent advances in mesoscopic imaging methods could allow resolution of larger cell populations and a direct linking of map and cellular levels in the same subjects (Cardin et al., 2020).

### Spatiotemporal features of a cortex-wide CSD in response to focal M1 injury

A prominent feature of our results was the presence of a large amplitude, slow moving wave of increased calcium signal activation that propagated across the cortex following M1-CCI, consistent with the well-described cortical spreading depression, also known as cortical spreading depolarization (CSD) (Dreier et al., 2011). CSD is initiated by mechanical trauma to neural tissue at the injury site, causing elevation in extracellular K+, contributing to propagation of the CSD via synaptic and non-synaptic depolarization mechanisms (Lauritzen et al., 2011). CSD typically occurs as a two-stage process: a large amplitude depolarization followed by a depression in overall activity (Bogdanov et al., 2016; Lauritzen et al., 2011; Enger et al., 2015; Charles and Brennan, 2009). The depolarization phase, initially causing neuronal excitation, can also suppress spiking activity through depolarization block, arising from inactivation of voltage-gated Na+ channels at depolarized membrane potentials. Calcium imaging directly captures the depolarization phase of the CSD, as GECIs are optimized to report depolarization and spiking activity. It is possible that calcium signals remain elevated even during depolarization block, since high-threshold calcium channels do not readily inactivate, and could diverge from CSD dynamics as recorded electrophysiologically. Decreased calcium signals from neural suppression are not as readily detectable compared to increases; however, we did also observe the CSD depression phase as a reduction of calcium signal fluorescence following the initial fluorescence increase. The most likely explanation for the calcium signal decrease is the suppression of sub- and suprathreshold cortical activity. This would be consistent with classic field potential or EEG recordings of TBI-triggered CSD, where the depression of cortical activity also dips below baseline levels. We did not observe such depressions in calcium signals in same-length recordings from uninjured mice, suggesting that these calcium signal decreases were not due to photobleaching. Thus, the depression phase of the CSD likely corresponds to the depression that we observed in the calcium signal. Overall, the wave velocity that we observed with widefield calcium imaging, its large amplitude, the secondary suppression, and its wide spatial extent, are consistent with the traditionally described CSD (Ayata and Lauritzen, 2015; Dreier et al., 2017). Our data add to a growing literature using imaging methods in unanesthetized mice to capture cortex-wide pathological events, such as CSD. Two recent studies imaged CSD after focal cortical application of KCl (Enger et al., 2017; Monai et al., 2021), while Cramer et al. (2023) performed widefield calcium imaging after repeated CCI application. Our experiments fill in the gap of imaging during CCI induction; similar to the KCl model, we are able to resolve the immediate effects as well as track the long-term effects in subsequent sessions. Our work emphasizes that imaging in transgenic reporter mice is well-suited to measure the spatiotemporal features of CSD in TBI especially during the application of injury, and for investigating long-term changes in sensory responses and functional connectivity over expanded spatiotemporal scales and with cell type specificity (Monai et al., 2021; Cramer et al., 2023) including neurons and astrocytes (Enger et al., 2017).

### Cellular changes

In our two-photon calcium imaging experiments in S1 barrel cortex, properties of the M1-CCI-evoked CSD were consistent with our widefield measures in terms of the large amplitude, slowly moving signal, followed by a global fluorescence decrease. The CSD wavefront appeared as a spreading neuropil signal that coincided with fluorescence increases in L2/3 neuronal somata. These data address the issue of involvement of neuronal somata in CSD (Charles and Brennan, 2009). It has been proposed that neuronal dendrites are particularly susceptible to CSD and are perhaps even exclusively involved in propagation of CSD. While our neuropil calcium signal includes contributions from axons and dendrites, we clearly observed somatic activation coincident or even preceding the neuropil signal at the CSD wavefront (Fig. 5). If dendritic preceded somatic activity, this occurred on a time scale faster than our 15 Hz frame rate. Thus, our data rule out the scenario that dendrites are active independent of cell bodies, at least in L2/3 of S1. However, we do not know the extent to which somatic calcium signals reflect spiking; it is possible that dendrites drive strong somatic depolarization and transient burst firing before depolarization block prevents further action potentials. Further experiments combining calcium imaging and electrophysiological recordings would be needed to resolve this issue.

After the CSD passed through S1, neuronal calcium signals remained elevated for several seconds, with sparse individual L2/3 neurons showing reactivation over the next minutes. This result suggests that subsets of excitatory L2/3 neurons show differences in susceptibility to TBI. One possibility is that these neurons comprise the M1-projecting, rather than S2-projecting, subpopulation (Chen et al., 2013), which had their axons damaged by the M1-CCI. Future experiments will need to determine whether these hyperactive neurons later become silent or remain hyperexcitable after injury. Chronic cellular-resolution imaging will be essential for tracking individual neurons from early post-TBI to long-term time points.

### Network changes

Network analysis of widefield calcium imaging data indicated an overall decrease in functional connectivity at early (20 and 60 min) post-TBI time points, with recovery by 24 h, consistent with recent imaging or electrophysiology studies measuring changes in cortical function after TBI (Bottom-Tanzer et al., 2024; Frankowski et al., 2021). The transient changes in neural activity that we observed paralleled the behavioral effects in our mild/moderate TBI protocol. Interestingly, recent work observed persistent disruptions in cortical network activity using bilateral calcium imaging after repeated mild TBI (Cramer et al., 2023) but without overt alterations in behavior. Stronger TBI paradigms that impair behavior, including stroke models, induce more dramatic changes in network activity, as measured with widefield calcium imaging or fMRI (Cramer et al., 2019; Harris et al., 2016). In the present study, the precise relationship between altered cortical activity and behavior remains unclear because we did not acquire detailed behavioral data during imaging (only afterward in the home cage). Thus we cannot rule out the possibility that the altered cortical activity we observed are due to TBI-induced changes in behavior rather than direct effects on cortical function. Using network analysis of neural responses to TBI has potential for comparison across imaging modalities (Aerts et al., 2016) and comparison of preclinical to clinical data (Pandit et al., 2013; Han et al., 2016). Application of frequency-resolved network analysis (Salsabilian et al., 2022) combined with neural activity sensors with increased temporal resolution (Yang and St-Pierre, 2016) and detailed behavioral analysis during imaging (Syeda et al., 2024; Mathis et al., 2018) would be a powerful advance.

### Sensory-evoked changes and relation to network changes

Sensory maps in S1 barrel cortex, in response to whisker deflection, increased in activation area at early post-TBI time points without an obvious change in amplitude. This result seems contradictory to the decreased number of active neurons we observed in two-photon recordings. One possibility is that deeper layer (L5) neurons, whose dendrites might contribute to the widefield signal, become hyperexcitable (Ding et al., 2011) while L2/3 neuron activity is reduced. Another possibility is that inhibitory neurons are suppressed, allowing a larger spread of sensory-evoked cortical activity (Isaacson and Scanziani, 2011). Since GECI was restricted to excitatory neurons in our Thy1 transgenic mouse line, our data do not address this possibility, and future experiments would be necessary to resolve TBI-induced changes in inhibitory neurons.

It is interesting to consider that the increase in the area of sensory-evoked S1 cortical maps occurred during the same post-CCI timeframe as the decrease in network connectivity. Two-photon cellular imaging data during this same timeframe suggested that the number of active S1 cells decreased, with a trend toward an increased number of calcium signals in these cells (see Fig. 8). These results suggest that the decrease in functional connectivity does not have a straightforward relationship with the underlying cellular activity. Rather, the relationship is more complex, possibly involving changes within populations of cortical neurons whereby subsets of neurons increase their activity and others decrease their activity, without a change in overall activity levels. Further studies will be necessary to determine the relationship between changes in functional connectivity with map- and cellular-level changes, possibly investigating homeostatic changes in both excitatory and inhibitory cortical networks.

### Limitations of the study

Our study had several limitations. First, while it would be interesting to understand the effects of TBI directly at the M1 injury site, we were not able to do this using our imaging methods because of local bleeding induced by the injury, which blocks optical access to the underlying tissue. Second, while our widefield imaging method allowed us to capture signals from an approximately 5×5 mm field of view, large enough to resolve many cortical areas from visual to motor cortex, larger fields of view would be desirable. Advances in cranial window implants spanning both cortical hemispheres as well as optical access to deep brain structures (Lee et al., 2020), along with improvements in widefield imaging technology, will be important for capturing the brain’s response to TBI across a larger spatial scale than we were able to do in the present study. Third, it is yet to be determined how results from rodent studies, such as ours, will translate to TBI of the human brain. Factors such as differences in TBI-induced edema, as well as factors related to morphological, cellular and circuit function could be important for clinical translatability. Finally, our study aimed largely to describe the effects of focal TBI on cortical function using a multiscale imaging approach, rather than investigate the detailed underlying mechanisms. While this may be seen as a limitation, we note that descriptive studies have been essential for identifying key problems and for providing a basis to guide further mechanistic work.

## Conclusions

Using widefield and two-photon calcium imaging in unanesthetized mice, we investigated the response to focal TBI across spatial and temporal scales. After the CSD induced by M1-CCI, we observed transient expansion of S1 sensory maps and decreased regional functional connectivity. Cellular imaging revealed subpopulations of L2/3 neurons in S1 with divergent responses to M1-CCI, suggesting unique susceptibility to injury. Overall, our results encourage a multiscale approach for investigating neural circuit adaptations to TBI, and emphasize that such experiments can be done in awake, head-restrained mice, avoiding confounding factors due to anesthesia.

## Acknowledgements

Funding provided by National Institutes of Health (NIH-R01NS094450) and National Science Foundation (IOS-1845355) grants to D.J.M., and NJ Commission on Brain Injury Research grant (CBIR161RG032) to D.J.M, L.N. and J.A. We thank Dr. Alex J. Yonk for expert help with graphics and figure layout.

## Author Contributions

Conceived and designed the research strategy: YB, DJM

Performed mouse husbandry and surgical procedures: YB

Performed imaging experiments: YB

Analyzed imaging experiments: YB, TJV, SS

Developed analytic tools and contributed reagents: TJV, SS, NF, JA, LN

Performed and analyzed behavior experiments: YB, AS, JF

Performed and analyzed histology experiments: YB, ST

Prepared the figures: YB, TJV, DJM

Wrote and edited the manuscript: YB, DJM with contributions from all authors.

Acquired funding: JA, LN, DJM

Supervised the project: DJM

The experimental work was performed in the Margolis laboratory at Rutgers. All authors approved the final version of the manuscript. All authors agree to be accountable for all aspects of the work in ensuring that questions related to the accuracy or integrity of any part of the work are appropriately investigated and resolved. All persons designated as authors qualify for authorship, and all those who qualify for authorship are listed.

## Data availability

Data and code are available at https://github.com/margolislab/Bibineyshvili_2024.

